# Inferring demographic and selective histories from population genomic data using a two-step approach in species with coding-sparse genomes: an application to human data

**DOI:** 10.1101/2024.09.19.613979

**Authors:** Vivak Soni, Jeffrey D. Jensen

## Abstract

The demographic history of a population, and the distribution of fitness effects (DFE) of newly arising mutations in functional genomic regions, are fundamental factors dictating both genetic variation and evolutionary trajectories. Although both demographic and DFE inference has been performed extensively in humans, these approaches have generally either been limited to simple demographic models involving a single population, or, where a complex population history has been inferred, without accounting for the potentially confounding effects of selection at linked sites. Taking advantage of the coding-sparse nature of the genome, we propose a 2-step approach in which coalescent simulations are first used to infer a complex multi-population demographic model, utilizing large non-functional regions that are likely free from the effects of background selection. We then use forward-in-time simulations to perform DFE inference in functional regions, conditional on the complex demography inferred and utilizing expected background selection effects in the estimation procedure. Throughout, recombination and mutation rate maps were used to account for the underlying empirical rate heterogeneity across the human genome. Importantly, within this framework it is possible to utilize and fit multiple aspects of the data, and this inference scheme represents a generalized approach for such large-scale inference in species with coding-sparse genomes.

## Introduction

Genetic variation is a fundamental concern of population genetics. Prior to the advent of next-generation sequencing, the dominant debate within the field was centered on whether levels of genetic variation were expected to be minimal or substantial (known as the *classical*/*balanced* debate; see Lewontin 1987; Crow 1987). Selection was assumed as the dominant process in both cases, be it purifying selection depressing levels of variation, or balancing selection maintaining polymorphism (Dobzhansky 1955). Despite molecular evidence confirming plentiful levels of genetic variation, Motoo Kimura’s Neutral Theory of Molecular Evolution (Kimura 1968, 1983) instead posited that observed variation was largely a consequence of genetic drift; that is, of neutral alleles segregating in the process of drifting towards fixation or loss. This hypothesis – that neutral rather than selective processes can explain the majority of observed variation – has since been largely corroborated (as reviewed in Jensen et al. 2019).

However, quantifying the precise roles of selective and neutral processes in shaping observed levels of variation – and disentangling their individual effects - remains an ongoing challenge due to the similar manners in which multiple evolutionary processes affect patterns of variation. One notable example is the extent to which neutral population growth, background selection (BGS; Charlesworth et al. 1993), and recurrent selective sweeps (Maynard Smith and Haigh 1974) can all skew the site frequency spectrum (SFS, the distribution of allele frequencies) toward rare alleles (Kim 2006; Jensen et al. 2007; Nicolaisen and Desai 2012, 2013; Ewing and Jensen 2016; Johri et al. 2021; Soni et al. 2023; and see review of Charlesworth and Jensen 2021, 2024). The effects of these processes are further modified by genomic heterogeneity in mutation and recombination rates in often complex ways (Soni et al. 2024b). Therefore, if one wishes to quantify the strength and frequency of rare and episodic processes such as positive selection, one must first construct an evolutionarily appropriate baseline model that accounts for the effects of constantly occurring processes including genetic drift as modulated by historical population size changes, as well as the effects of purifying selection and BGS resulting from the removal of deleterious mutations (Bank et al. 2014; Johri et al. 2022a), all whilst accounting for underlying mutation and recombination rate variation. Failure to account for these processes is likely to lead to misinference, particularly in light of the fact that many commonly studied populations and species are thought to have experienced not only population growth, but also recent and severe population bottlenecks [e.g. humans (Gutenkunst et al. 2009; Gravel et al. 2011; Excoffier et al. 2013), non-human primates (Terbot et al. 2024; Soni et al. 2024c) and Drosophila melanogaster (Li and Stephan 2006), as well as a variety of human pathogens (Irwin et al. 2016; Sackman et al. 2019; Jensen 2021; Morales-Arce et al. 2021)], a demographic history that is itself often strongly confounded with selective sweeps (Barton 1998; Poh et al. 2014; Matuszewski et al. 2018; Harris and Jensen 2020; Charlesworth and Jensen 2022; Jensen 2023).

Constructing an evolutionarily appropriate baseline model for a given population will therefore require inferring both a demographic history as well as the distribution of fitness effects (DFE) of new mutations. However, because population history can confound DFE inference, it is necessary to correct for the demographic history of the population in question (Eyre-Walker and Keightley 2007; Boyko et al. 2008). The most commonly used class of approaches are based on a framework in which demographic inference is performed on putatively neutral sites, before utilizing that demographic history for DFE inference on functional sites (Eyre-Walker and Keightley 2007; Boyko et al. 2008; Galtier 2016; Tataru and Bataillon 2020; and see review of Johri et al. 2022b). Eyre-Walker and Keightley (2007) obtained the first computationally inferred DFE estimates using this approach, and further work incorporated a beneficial class of mutations into the inferred DFE (Boyko et al. 2008; Eyre-Walker and Keightley 2009; Schneider et al. 2011; Galtier 2016).

Notably, this type of 2-step approach is often performed on functional regions under the assumption that all sites are independent and unlinked, and that synonymous sites are selectively neutral. However, these synonymous sites are likely experiencing BGS effects (Charlesworth et al. 1993) due to linkage with directly selected and adjacent non-synonymous sites, resulting in a skew in the SFS and thus mis-inference; in particular, these BGS effects are often misinterpreted as population growth (Ewing and Jensen 2014; Johri et al. 2021; and see review of Johri et al. 2022b). More generally speaking, there is indeed substantial evidence that the effects of selection at linked sites may be widespread across the genomes of many commonly studies species (see reviews of Cutter and Payseur 2013; Charlesworth and Jensen 2021). Although recent work has shown that DFE inference is relatively robust to the biasing effects of selection at linked sites (Kim et al. 2017; Huang et al. 2021), that is not the case for demographic inference (Messer and Petrov 2013; Nicolaisen and Desai 2013; Ewing and Jensen 2016; Schrider et al. 2016; Johri et al. 2021). It is also noteworthy that these 2-step approaches are generally constrained to relatively simple population histories utilizing a two-epoch model (Williamson et al. 2005; Keightley and Eyre-Walker 2007; Kousanthanas and Keightley 2013).

The second class of methods involve using forward-in-time simulations (e.g., in SLiM; Haller and Messer 2023) to jointly and simultaneously infer population history with the DFE in an approximate Bayesian (ABC) framework (see Beaumont et al. 2002), as proposed by Johri et al. (2020). Within this simultaneous inference scheme, it is neither necessary to assume *a priori* the neutrality of synonymous sites, nor is it necessary to assume independence amongst sites; as such, background selection can be directly modelled and incorporated. While 2-step methods commonly infer a continuous distribution for the DFE, this ABC framework infers a number of discrete DFE categories for various ranges of 2*N_e_s*, the population-scaled selection coefficient, where *N­_e_* is the effective population size and *s* is the strength of selection acting on new mutations within the DFE category of interest. The main drawback of such methods is that they are computationally expensive given the large parameter space that must be explored when jointly inferring both demographic and DFE parameters. As such, the inferred demographic models have thus far been limited to single-step size changes in which the ancestral and current population sizes, as well as the timing of size change, are inferred (Johri et al. 2020, 2023). Importantly however, in coding-dense and/or non-recombining species in which sufficiently neutral, unlinked genomic regions may not exist in the genome (thus precluding the needed neutral demographic inference underlying 2-step approaches), this simultaneous inference framework remains the only viable approach (e.g., Howell et al. 2023; Terbot et al. 2023a,b; Soni et al. 2024a).

It thus stands as an outstanding evolutionary inference question of how best to accurately infer a necessarily complex and realistic demographic model, along with a realistic DFE governing functional genomic regions, all whilst accounting for the variety of discussed potential biases. Here we have investigated a modified 2-step approach applied to human populations, in which the population history was inferred using non-functional regions sufficiently distant from functional sites in order to avoid BGS effects, DFE inference was then performed on exonic regions accounting for BGS effects and conditional on the demographic history inferred in Step 1, and mutation and recombination rate maps were utilized to account for the modulating effects of this underlying heterogeneity. By inferring these parameters separately, a more biologically realistic population history was possible accounting for the complexities of population size change, structure, and migration patterns in these studied human populations, while the utilization of these distant non-functional regions allowed for the reduction or elimination of the biasing effects of BGS on demographic inference. Whilst a number of coalescent and diffusion approximation-based approaches would be easily incorporated into our framework (e.g., Gutenkunst et al. 2009; Excoffier et al. 2013; Jouganous et al. 2017; Wang et al. 2020), this approach – like the ABC approach of Johri et al. (2020, 2022a) - has the benefit of utilizing various aspects of population genomic data, including the SFS, associations between variants (linkage disequilibrium, LD), and population differentiation.

As human populations have naturally been highly studied, with numerous published demographic models, we here provide an optimized and well-fitting 4-population demographic model for the Out-of-Africa (OOA) expansion. Conditional on this model, we additionally optimized a DFE using genic regions, fitting both levels and patterns of polymorphism and divergence, and finding consistency with the recent DFE estimates of Johri et al. (2023). Finally, we have evaluated the degree to which positively selected mutations may be identifiable within the context of this fit model. This work thus provides a valuable and improved framework for evolutionary inference in coding-sparse genomes, and for the construction of evolutionary baseline models in such species.

## Methods and Materials

### Data

This study was based on the GRCh37 human reference genome, with SNP data and accessibility masks obtained from 1000 genomes variant call format and bed files, respectively (The 1000 Genomes Project Consortium 2015). The data was split into continental populations, informed by levels of admixture, as determined by The 1000 Genomes Project Consortium (2015). The total number of samples from each of the four considered populations were: African – 99; European – 502; East Asian – 104; South Asian – 489. We obtained recombination and mutation rate maps from Halldorsson et al. (2019) and Francioli et al. (2015), respectively, gene annotations from NCBI (Sayers et al. 2022), ancestral sequences from the six-way EPO alignments available from Ensembl (Flicek et al. 2014; Cunningham et al. 2022), and we identified conserved elements via the 100-way PhastCons score (Siepel et al. 2005; Pollard et al. 2010). See Supplementary Table S1 for links to all downloaded data.

### Selecting non-functional regions for demographic inference

For demographic inference we identified non-functional regions of the human genome that were at a distance of at least 10kb from the nearest functional region (as per the NCBI GFF file [Sayers et al. 2022]). We then masked these regions using both strict accessibility masking (The 1000 Genomes Project Consortium 2015) and conserved element masks (i.e., with a phastCons score > 0 [Siepel et al. 2005; Pollard et al. 2010], in order to remove sites potentially experiencing purifying selection and generating background selection effects (e.g., binding sites (Simkin et al. 2014))). Across each region, we calculated mean recombination and mutation rates, with any regions lacking this information being removed. Finally, we set a minimum length threshold of 15kb to ensure that regions were long enough to reliably calculate summary statistics. Following these steps, we were left with a total of 146 non-functional regions. Finally, we used the B maps of McVicker et al. (2009) to compare the distribution of B values (i.e., the estimated reduction in diversity attributed to BGS by McVicker et al. (2009)) to the distribution of our non-functional regions. For this analysis we lifted over B map coordinates from the hg18 human genome assembly to the GRCh37 assembly using the UCSC liftover tool (Karolchik et al. 2003). Supplementary Figure S1 provides plots of this comparison, as well as the distributions of region lengths, SNPs, and mutation and recombination rates for our set of curated non-functional regions.

### Selecting exons for DFE inference

We used the set of exons curated by Johri et al. (2023), although our focus was on the exonic regions only, as opposed to the exons and the neighbouring intergenic regions. Because we used different recombination and mutation rate maps (as described in the data section above), we recalculated mean rates across the 465 exonic regions, removing regions for which we did not have rate information, leaving a total of 397 exonic regions.

### Calculating empirical summary statistics

We calculated summary statistics for each population sampled using the python library for libsequence, Pylibseq v0.2.3 (Thornton 2003), except for *F_ST_* which was estimated using scikit-allel (Miles et al. 2024). The number of segregating sites and *F_ST_* were calculated per site, whilst Tajima’s *D* (Tajima 1989) and mean *r^2^* were calculated over 10kb windows for each non-functional region, and per region for each exon.

Exonic divergence was calculated based on the number of fixed differences between the reference and ancestral sequences, with polymorphic sites masked, relative to total region size.

### Calculating summary statistics from simulated data

We calculated summary statistics from simulated data in a manner that replicated the empirical data, using the same software as described above. Thus, sites that had been masked in the empirical data were also masked in the simulated data prior to calculating summary statistics.

Exonic divergence was calculated as the number of fixations post-burn-in from forward-in-time simulations (see the ’Simulating human population history with selection using SLiM’ section).

We calculated the mean and standard deviation for each region across its respective 100 replicates. For plotting purposes, we plotted the mean of all regions as the data point, and the mean of the standard deviations across all regions as the confidence intervals.

### Simulating human population history using Msprime

Step 1 in our 2-step inference framework was the inference of population history. We simulated human demography using the coalescent simulator Msprime (Baumdicker et al. 2022) for each of our 146 non-functional regions, with region specific mutation and recombination rates. Our demographic model was comprised of 5 populations (four sampled populations: African, European, South Asian and East Asian; as well as the unsampled ancestral Eurasian population) and 25 parameters. Parameter ranges were taken from the human demographic inference literature, with midpoints of all ranges used as the initial starting parameterization. A generation time of 26.9 years was used to appropriately scale simulations (Wang et al. 2023). For details of the demographic model see Figure 1 and Table S2. 100 replicates were simulated for each of the 146 non-functional regions, with a single mutation and recombination rate per region, calculated as the average across the region from the Francioli et al. (2015) mutation rate map and the Halldorsson et al. (2019) recombination rate map (see Supplementary Figure S1 for distributions of region lengths, mutation rates and recombination rates across curated regions). Parameters were optimized to the data using *F_ST_*, the number of segregating sites, Tajima’s *D* (Tajima 1989) and mean *r^2^*, across all four populations. Demographic inference plots (e.g., Figure 1) were produced using Demes software (Gower et al. 2022).

**Figure 1:**
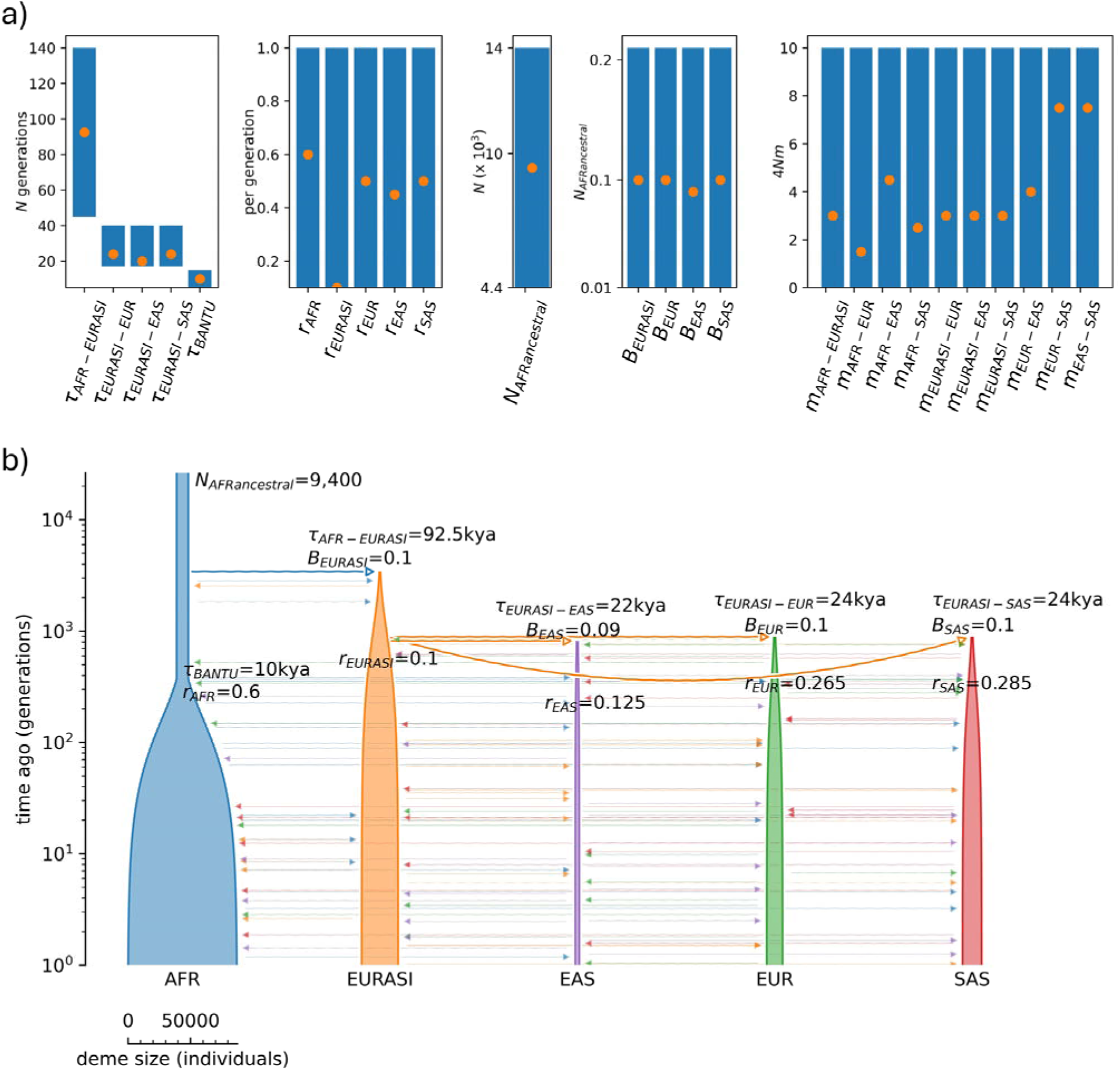
Demographic model representing the Out-of-Africa expansion. a) Parameter ranges for all 25 parameters (represented by the blue bars on the plots). Orange dots indicate the best fitting parameter values identified. b) Plot of demographic model with the best fitting parameter values. Population key: AFRancestral = initial ancestral African population; AFR = African population; EURASI = unsampled Eurasian population; EUR = European population; EAS = East Asian population; SAS = South Asian population. Parameter key: *τ* = time of splits between specified populations (with *τ_BANTU_* representing the time of start of the Bantu expansion in the African population); *r* = growth parameter; *N* = population size; *B* = bottleneck severity; *m* = migration rate. Demographic model graphic generated using Demes software (Gower et al. 2022).

### Simulating human population history with selection using SLiM

For Step 2, we simulated the inferred population history from Step 1 using the forward-in-time simulator SLiM (v4.0.1 [Haller and Messer 2023]), for our 397 exonic regions, with region specific mutation and recombination rates. We simulated to the human-chimpanzee split (12mya; Moorjani et al. 2016). Thus, the simulations considered 12 million years (446,100 generations) before starting the 10*N* generation burn-in period. Exonic mutations were drawn from a DFE comprised of 4 fixed classes (following Johri *et al*. 2020), with frequencies denoted by *fi*: *f*_0_ with 0 ≤ 2*N_AFRancestral_ s* < 1 (i.e., effectively neutral mutations), *f*_1_ with *1* ≤ 2*N_AFRancestral_ s* < 10 (i.e., weakly deleterious mutations), *f*_2_ with 10≤ 2*N_AFRancestral_ s* < 100 (i.e., moderately deleterious mutations), and *f*_3_ with 100 ≤ 2*N_AFRancestral_ s* (i.e., strongly deleterious mutations), where 2*N_AFRancestral_* is the initial African population size and *s* is the reduction in fitness of the mutant homozygote relative to the wild-type. We initially simulated the DFE from Johri et al. (2023) - comprised of neutral and deleterious mutations - which fit the empirical data well.

### Simulating selective sweeps

#### Recurrent

We simulated recurrent selective sweeps by adding a beneficial DFE category for our 397 exonic regions. We simulated three different beneficial rates (0.1%, 1%, and 10% of new mutations), with the effectively neutral DFE category (*f_0_*) reduced to account for the addition of the beneficial category. Three different beneficial classes were separately simulated: 1 ≤ 2*N_AFRancestral_ s_b_* < 10; 10 ≤ 2*N_AFRancestral_ s_b_* < 100 and 100 ≤ 2*N_AFRancestral_ s_b_* < 1,000, where *s_b_* is the increase in mutant homozygote fitness relative to the wild-type.

#### Individual

To simulate a single hard selective sweep, we ran our inferred demographic model with the inferred DFE, with three different scenarios for introducing a beneficial mutation: model 1) the beneficial mutation was introduced into the African population immediately after burn-in; model 2) the beneficial mutation was introduced into the ancestral Eurasian population immediately after splitting from the African population; and model 3) the beneficial mutation was introduced into the European population immediately after splitting from the Eurasian population. In model 1, simulations were terminated and restarted if the beneficial mutation did not fix in all 4 sampled populations. In model 2, simulations were terminated and restarted if the hard sweep did not fix in the European, East Asian and South Asian populations. Finally, in model 3 simulations were terminated and restarted if the hard sweep did not fix in the European population. For each scenario, two different strengths of selection were simulated: 2*N_e_s_b_*= 1,000 and 10,000, where *N_e_* is the ancestral African population size (*N_AFRancestral_*) and *s_b_* is the beneficial selection coefficient.

For these simulations, we utilized the chromosomal structure of Soni and Jensen (2024), with functional regions comprised of 9 exons (each of size 1,317bp) and 8 introns (each of size 1,520bp), separated by intergenic regions (each of size 4,322bp) [The 1000 Genomes Project Consortium 2015]. The number of exons and introns per functional region were taken from Sakharkar et al. (2004). The chromosomal region contained 7 functional regions in total, resulting in a total simulated region length of 198,345bp.

Variable mutation and recombination rates were drawn from a uniform distribution such that the mean recombination rate across the simulated region for each replicate was equal to the Kong et al. (2010) mean, and the mean mutation rate across the simulated region for each replicate was equal to the Kessler et al. (2020) mean.

### Sweep inference with SweepFinder2

We performed selective sweep inference by running SweepFinder2 (DeGiorgio et al. 2016) on each simulated replicate of each exonic region from our hard sweep simulations. Allele frequency files were generated for each replicate, following Huber et al.’s (2016) recommendation of including only polymorphic and substitution data. Inference was performed at each SNP via a grid file, following Nielsen et al. (2005). The background SFS was taken from the sweep-free simulations inferred in this study. The following command line was used for inference: SweepFinder2 -lru GridFile FreqFile SpectFile RecFile OutFile

### Sweep inference with H12

We ran the H12 method of Garud et al. (2015) on each simulated replicate of each exonic region from our hard sweep simulations, using a custom python script. H12 was estimated over 1kb, 2kb, 5kb, 10kb, 20kb, and 40kb windows at each SNP, with the SNP at the center of each window.

For both SweepFinder2 and H12 inference, we calculated true- and false-positive rates based on the inference values at each site, generating ROC curves from this information.

### Results and Discussion

Our implemented 2-step approach to demographic and DFE inference involves inferring population history using non-functional regions that are at a sufficient distance from functional sites so as to reasonably ensure that they are not experiencing purifying or background selection effects. DFE inference is then performed on exonic regions in Step 2, conditional on the demographic history inferred in Step 1 and incorporating expected background selection effects. We have applied this approach to human population genomic data from the 1000 genomes project (The 1000 Genomes Project Consortium 2015), in order to better characterize the evolutionary parameters governing recent human history.

### Step 1: Demographic inference on non-functional regions

In order to avoid the biasing effects of purifying selection and BGS, we performed demographic inference on our curated set of 146 non-functional regions, with mean recombination and mutation rates calculated for each region from the rate maps of Halldorsson et al. (2019) and Francioli et al. (2015), respectively. For details of the data curation steps, please see the Methods section. While one would typically begin with an evaluation of numerous demographic models and topologies in less well-characterized species (see Beaumont et al. 2002; Johri et al. 2020), given the considerable literature on human demographic history (e.g., Gutenkunst et al. 2009; Gravel et al. 2011; Schiffels and Durbin 2014; Terhorst et al. 2017; Hu et al. 2023), and inferred levels of admixture in The 1000 Genomes dataset (The 1000 Genomes Project Consortium 2015), we began with a model of the Out-Of-Africa (OOA) colonization in which the ancestral Eurasian population splits from the African population, followed by the European, South Asian and East Asian populations dispersing from the ancestral Eurasian population, along with the Bantu expansion in the African population. Thus, our demographic model was comprised of 5 populations (African, ancestral Eurasian, European, South Asian and East Asian, of which all but the ancestral Eurasian population were sampled) and 25 parameters that capture population sizes, bottleneck severities, growth rates, timings of each event, and migration rates between populations. Parameter ranges were drawn from the extensive literature on human population history (Mellars 2006; Gutenkunst et al. 2009; Gravel et al. 2011; Tennessen et al. 2012; Terhorst et al. 2017). Figure 1a provides the parameter ranges for our model, and see Methods section for further details.

We simulated 100 replicates for each of our 146 non-functional regions using the coalescent simulator MSprime (Baumdicker et al. 2022) with region-specific mutation and recombination rates, initially starting with midpoint values for each of our parameters (see Figure 1a). For each replicate we estimated four summary statistics for each population (or pairs of populations): the number of segregating sites, Tajima’s *D* (Tajima 1989), mean *r^2^*, and *F_ST_*, giving us a total of 18 summary statistics. Fitting these four statistics enabled us to account for multiple aspects of the data including levels of diversity, the SFS, LD and population structure. Figure 1a provides the optimized fit of each parameter within the context of previously published parameter ranges, and Figure 1b the total inferred demographic model. As shown in Figure 2, the summary statistics resulting from this demographic model well fit observed empirical data.

**Figure 2:**
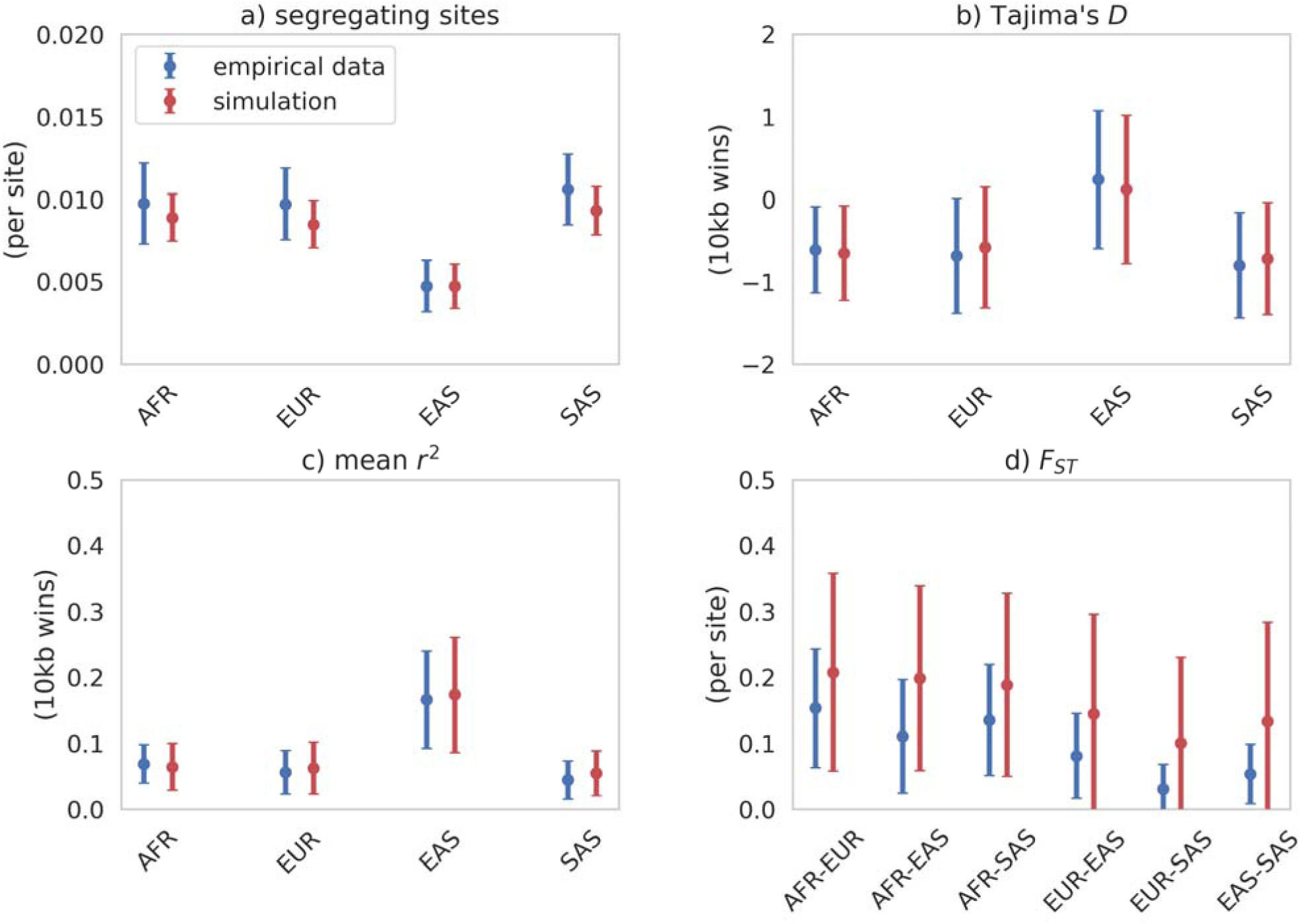
Summary statistics calculated from putatively neutral non-functional regions from population samples for empirical (blue) data, compared to simulated (red) data under the best-fitting demographic model. Means and standard deviations were calculated for 100 replicates. Data points represent the mean across regions, while bars represent the mean of the standard deviations across all regions.

It is notable that the African population in our model is larger than the African populations in the Gutenkunst et al. (2009) and Gravel et al. (2011) best-fitting models. There are two likely contributing factors. Firstly, these previous studies fit the model to the SFS, whereas we have here fit multiple diverse summaries of the data. Secondly, these previous studies modeled the African population with a fixed size that undergoes a single instantaneous expansion. Here we modelled the recent Bantu expansion, and thus our final African population size was notably larger, though our final African population size of 87,594 falls within the range of previous estimates (Schiffels and Durbin 2014; Terhorst et al. 2017; Johri et al. 2023). Finally, it is worth noting that numerous other coalescent and diffusion approximation-based approaches have been used to infer the OOA model of human population history (Gutenkunst et al. 2009; Gravel et al. 2011; Excoffier et al. 2013; Jouganous et al. 2017; Wang et al. 2020). These studies have masked genic regions to avoid the biasing effects of selection. However, BGS can still affect demographic inference if not accounted for; nonetheless, our parameter estimates fall within previously inferred ranges, confirming the modest nature of BGS effects in humans (Johri et al. 2021; Buffalo and Kern 2024).

In summary, by optimizing within previously published parameter ranges, we have identified a neutral demographic model that well explains multiple facets of the genomic data in distant non-coding regions.

### Step 2: DFE inference on functional regions

Given the strong fit of the neutral demographic model to the intergenic data, we next moved to Step 2: inference of the DFE using functional regions. We utilized the curated set of functional regions from Johri et al. (2023). After obtaining region-specific mutation and recombination rates we were left with a total of 397 functional regions. Unlike Johri et al. (2023) who simulated exons and their neighboring regions, we focused on the exons only (given that the model fit was consistent across both exons and adjacent regions in their study). First, we simulated our 397 functional regions under the demographic model inferred in Step 1, using the forward-in-time simulator SLiM (v4.0.1 [Haller and Messer 2023]). For the purpose of DFE inference, we simulated to the human-chimpanzee split time (12mya [Moorjani et al. 2016]) to allow us to compare empirical and simulated divergence, which is expected to be shaped by selection at functional sites. When simulating these functional regions under selective neutrality, we found that the fit to the empirical data was poorer than for the non-functional regions (Supplementary Figure S2); an expected result given the action of selection in these exonic regions. Next, we simulated under the Johri et al. (2023) DFE using our fit demographic model, and found a good fit of the simulated summary statistics to the empirical data (Figure 3). These results are encouraging given the differing approaches taken between the two studies: we here took the 2-step approach as described, whilst Johri et al. utilized a simultaneous inference scheme. Importantly however, both studies accounted for expected BGS effects, a relative rarity in DFE inference.

**Figure 3:**
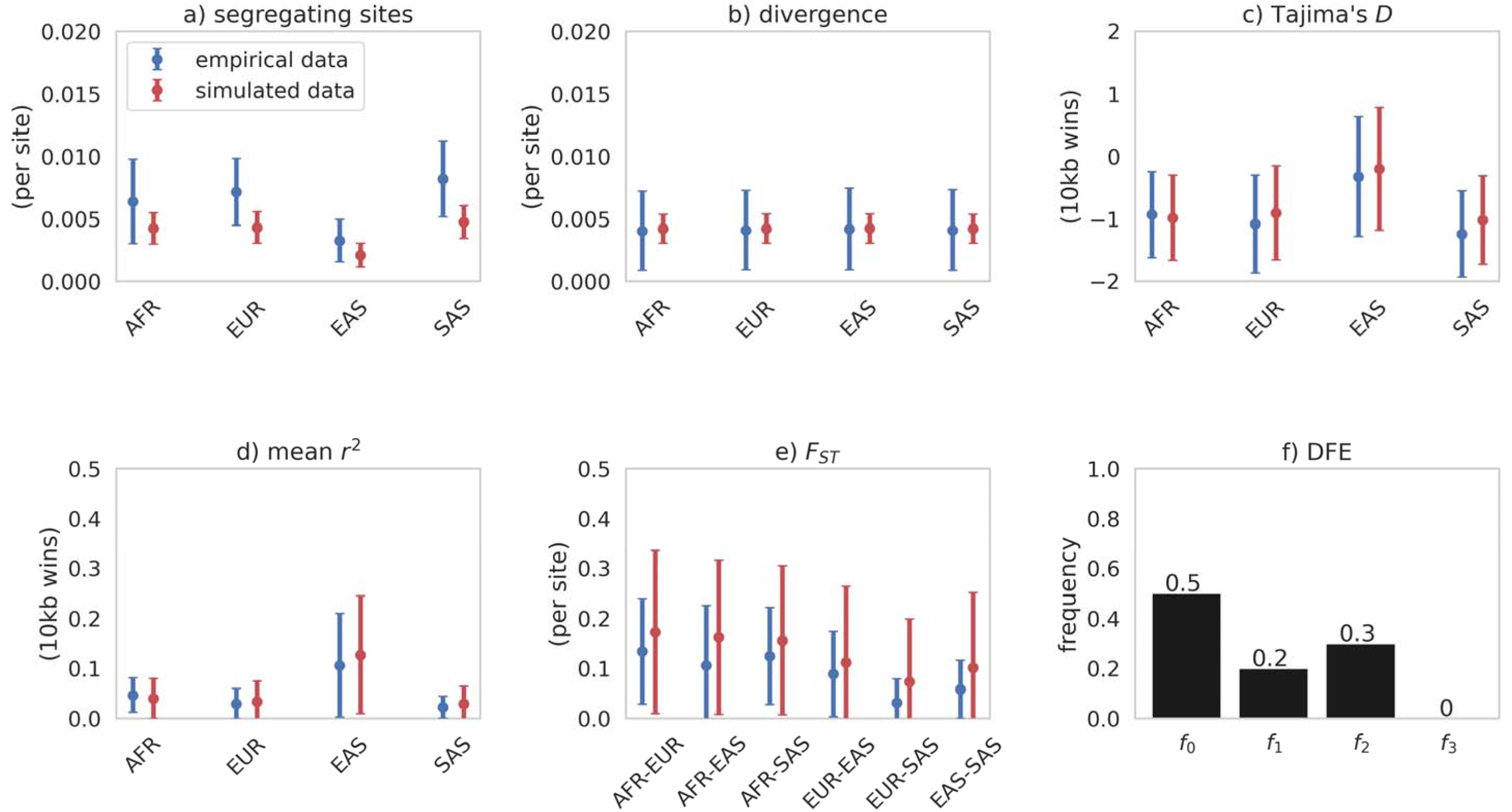
a) to e) Summary statistics calculated from functional regions from population samples for empirical (blue) data, compared to simulated (red) data under the best-fitting neutral demographic model with the addition of purifying and background selection modelled using the Johri et al. (2023) DFE (shown in panel f). Following this DFE, exonic mutations were drawn from a DFE comprised of 4 fixed classes with frequencies denoted by *f_i_*: *f_0_* with 0 ≤ 2*N_AFRancestral_ s* < 1 (i.e., effectively neutral mutations), *f_1_* with 1 ≤ 2*N_AFRancestral_ s* < 10 (i.e., weakly deleterious mutations), *f_2_* with 10 ≤ 2*N_AFRancestra_l s* < 100 (i.e., moderately deleterious mutations), and *f_3_* with 100 ≤ 2*N_AFRancestral_ s* (i.e., strongly deleterious mutations), where *N_AFRancestral_* was the initial population size and *s* the reduction in fitness of the mutant homozygote relative to wild-type. Means and standard deviations were calculated for 100 replicates. Data points represent the mean across regions, while bars represent the mean of the standard deviations across all regions.

Though the inclusion of population history, purifying and background selection effects, and mutation and recombination rate heterogeneity were alone sufficient to explain empirically observed data patterns, that does not necessarily imply the absence of beneficial mutations; rather, it suggests that this additional parameter is not needed in order to fit observed patterns of variation. While this observation is itself meaningful, it indeed raises the question of what rate of beneficial mutation may be consistent with the data but simply unidentifiable. In order to investigate this question, we added a beneficial DFE category to the Johri et al. (2023) DFE, in an attempt to understand what rate of input of beneficial mutations may be compatible with the observed levels of variation, the SFS, LD, divergence and *F_ST_*. Initially, we considered three beneficial DFE proportions, *f_bo_* = [0.1%, 1%, or 10% of newly arising mutations], with 1 ≤ 2*N_AFRancestral_ s_b_* < 10 (i.e., weakly beneficial mutations). Under this model, we correspondingly reduced *f_0_* - the proportion of effectively neutral mutations – in order to account for the addition of this beneficial DFE class. Supplementary Figures S3-S5 provide the fit of the summary statistics from these simulations to the observed data. At *f_b0_* = 0.1% or 1%, all summary statistics remain reasonably well fit - in other words, they are not significantly modified from the expectations in the absence of positive selection. However, divergence was notably increased relative to that observed at *f_b0_* = 10%, due to the greater rate of beneficial fixation.

Given that this beneficial mutation rate of 10% appears inconsistent with empirical divergence, we next examined *f_b0_* = 0.1% and 1% only, whilst increasing the population-scaled strength of selection to 10 ≤ 2*N_AFRancestral_ s_b_* < 100 (i.e., moderately beneficial mutations). Supplementary Figures S6 and S7 provide the fit of summary statistics from these simulations to the observed data. With this increased strength of selection, the modelled divergence only fit the empirical data at the lowest beneficial frequency, *f_B0_* = 0.1%. Finally, we increased the population-scaled strength of selection further to 100 ≤ 2*N_AFRancestral_ s_b_* < 1000 (i.e., strongly beneficial mutations), at *f_B0_* = 0.1%. Even at this low frequency, the resulting divergence was too high relative to the empirical data (see Supplementary Figure S8 for all summary statistics). It is notable that regardless of beneficial mutation frequency or strength of selection, the other summary statistics fit the data well - this owes to the relative waiting time between selective sweeps under these models; that is, selective sweeps are too old on average to strongly impact patterns of polymorphism (Jensen 2009), while being frequent enough to modify divergence over the 12mya time-scale.

Taken together, these results suggest that whilst the addition of a beneficial DFE class is not necessary to explain the patterns observed in the human population genomic data here considered, a modest input of weakly beneficial mutations and/or a low input of moderately beneficial mutations would remain consistent with the observed data.

### Evaluating power to detect selective sweeps within this human baseline model

Recurrent sweep models, such as the one studied above, involve a scenario in which beneficial mutations occur randomly across a chromosome according to a time-homogenous Poisson process at a per-generation rate (Kaplan et al. 1989; Wiehe and Stephan 1993; Stephan 1995; Pavlidis et al. 2010; Soni et al. 2023). Although this is a more realistic model of positive selection, in that the beneficial mutations underlying selective sweeps naturally occur at a per-generation rate - meaning that they are naturally associated with an average time since fixation - the more commonly studied model involves a single selective sweep in which fixation occurred immediately prior to sampling. As such, these models consider a best-case scenario for sweep detection, both in that sweeps are as recent as possible thus maximizing detectable polymorphism-based patterns (see review of Nielsen 2005), but also because it avoids the possibility of interference between positively selected mutations (i.e., Hill and Robertson 1966).

Furthermore, these models are often simulated on otherwise neutral backgrounds, which is additionally unrealistic in the sense that beneficial mutations occur in functional regions, which will be dominated by newly arising deleterious mutations. Thus, as a step towards biological reality, we here have modelled single selective sweeps within the context of our evolutionary baseline model, using our inferred demographic history, DFE, as well as mutation and recombination rate maps, thereby accounting for constantly-operating evolutionary processes in order to characterize the power to identify an episodic selective sweep (as described by Johri et al. 2022a).

Under this model, we simulated a large genomic region comprised of functional and non-functional regions in which a single hard selective sweep occurred in a functional element (see Methods section for more details about simulated chromosomal structure, as well as parameterizations). Sweep inference was conducted using two methods: the composite-likelihood ratio (CLR) SFS-based method, SweepFinder2 (DeGiorgio et al. 2016), and a haplotype-based approach, H12. Three different sweep models were simulated: 1) a beneficial mutation introduced into the ancestral African population immediately after simulation burn-in, with the fixed beneficial present in the sampled African, European, East Asian and South Asian populations; 2) a beneficial mutation introduced into the ancestral Eurasian population immediately after splitting from the ancestral African population, with the fixed beneficial present in the sampled European, East Asian and South Asian populations; and 3) a beneficial mutation introduced into the European population immediately after splitting from the Eurasian population, with the fixed beneficial present in the sampled European population. Figure 4 presents ROC plots, plotting the false positive rate (FPR) against the true positive rate (TPR) for inference on each model across 100 replicates with SweepFinder2 (with inference performed at each SNP) and H12 (with inference performed across 1kb windows, centered on each SNP; see Supplementary Figures S9-13 for additional window sizes).

**Figure 4:**
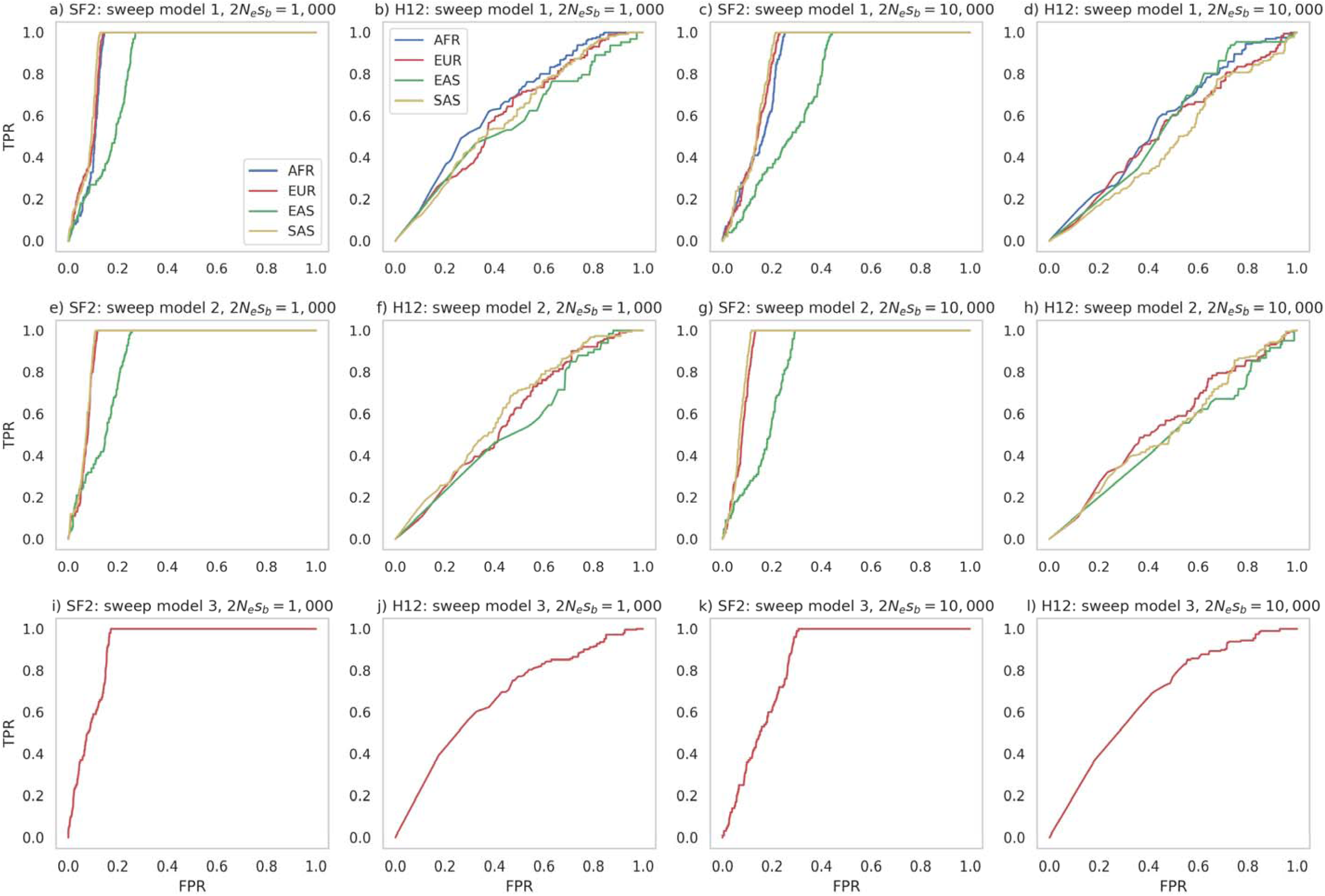
ROC curves, providing the change in true-positive rate (TPR) with false-positive rate (FPR), for sweep inference with SweepFinder2 (SF2) and the H12 statistic under the demographic model inferred in this study (see Figure 1) together with the Johri et al. (2023) DFE for functional regions, and variable mutation and recombination rates (see Methods section). Here, a single beneficial mutation was introduced into the population at three different time points and in three different populations: Model 1: the beneficial mutation was introduced into the ancestral African population immediately after the burn-in period, the beneficial fixation is present in all populations, and sweep inference was conducted on all sampled populations; Model 2: the beneficial mutation was introduced into the ancestral Eurasian population immediately upon splitting from the ancestral African population, the beneficial fixation is present in all non-African populations, and sweep inference was conducted on the European, East Asian and South Asian populations; Model 3: the beneficial mutation was introduced into the European population immediately upon splitting from the Eurasian population, the beneficial fixation is present in the European population, and sweep inference was conducted on this population only. For each model, two different strengths of selection were modelled: 2*N_e_s_b_*= 1,000 and 2*N_e_s_b_* = 10,000, where *N_e_* is the size of the ancestral African population and *s_b_* is the selection coefficient of the beneficial mutation. Inference with SweepFinder2 was performed on each SNP and substitution, whilst H12 inference was performed on each SNP over a 1kb window, with the SNP at the center of the window.

At the lowest strength of selection (2*N_e_s_b_* = 100), no beneficial mutations reached fixation by the sampling time (i.e., the present day) across the replicates. As such, Figure 4 presents ROC plots for 2*N_e_s_b_* values of 1,000 and 10,000 only. Although SweepFinder2 showed greater inference power than H12, there was limited power to detect selective sweeps for both approaches. While potentially appearing counter-intuitive, in some circumstances 2*N_e_s_b_* = 1,000 had greater power than 2*N_e_s_b_* = 10,000, as the fixations of the former were more recent given the longer sojourn time, and thus experienced less post-fixation decay in patterns of polymorphism (Kim and Stephan 2000; Soni et al. 2023). These results thus suggest that detectable selective sweeps would necessarily be the result of positive selection that was strong and recent enough to leave a detectable signature, consistent with previous work (Przeworski 2002; Kim and Stephan 2002; Jensen et al. 2007; Crisci et al. 2013). Moreover, the modest power under our baseline model is likely explained by the severe bottlenecks and expansions characterizing these populations, as the fundamental difficulty in distinguishing between population bottlenecks and selective sweeps has been previously demonstrated (Barton 1998; Jensen et al. 2005). These results suggest that caution is needed when performing genomic scans for selection in humans due to their complex recent demographic history, and likely supports previous assertions that strong selective sweeps have been rare in recent human history (Hernandez et al. 2011).

## Conclusions

In this study we have demonstrated the viability of a 2-step approach for inferring population history along with the DFE in coding-sparse genomes, such as that characterizing humans. This condition, together with being a recombining genome, is important for the existence and availability of non-functional regions sufficiently distant from functional sites so as to be free from the effects of purifying and background selection, as such regions are necessary for the accurate inference of population history. By contrast, organisms with genomes that are either coding dense or experience limited recombination may not have such regions in sufficient number, in which case demographic inference must be performed within the context of background selection effects. As these background selection effects will be dictated partially by the DFE in functional regions, this genomic architecture requires the joint and simultaneous inference of demographic and selective parameters - a situation that spans organisms ranging from *Drosophila* to many viruses (see review of Johri et al. 2022b). However, given the multiple jointly inferred parameters, the demographic histories under these joint inference schemes have been highly simplified in current implementations. Thus, this 2-step approach has a distinct advantage for coding-sparse genomes, in that previously developed and sophisticated neutral demographic inference approaches may be leveraged in Step 1 - such as that employed here estimating a 25-parameter human demographic model consisting of multiple population size changes, split times, and migration rates - allowing DFE inference to be focused upon in Step 2 conditional on that inferred history.

It is additionally important to consider the extent to which a consideration of these BGS effects matters for human demographic inference. Indeed, given the coding-sparseness of the genome, these effects are expected *a priori* to be limited, and that is fully consistent with the observation that our optimized demographic parameter values fall within previously published parameter ranges. However, apart from accounting for the effects of selection at linked sites, this approach also utilizes patterns of variation in addition to the site frequency spectrum (e.g., linkage disequilibrium and population-differentiation), which provide a further valuable ’sanity check’ on estimated models. This combination of factors has resulted in incrementally improved - but indeed improved - parameter estimates for the populations studied, as assessed by the fit between the estimated model and the empirical data. Thus, this proof-of-principle approach applied here to publicly-available human data will likely provide a highly relevant and informative inference framework for the analysis of future genomic resources in comparatively poorly-studied species with a similar genomic architecture (e.g., non-human primates).

