## Supplementary for "Inferring demographic and selective histories from population genomic data using a two-step approach in species with coding-sparse genomes: an application to human data"

**Table S1**

| **Data** | **Link** |
|---|---|
| GRCh37 Gene annotations | https://ftp.ncbi.nlm.nih.gov/refseq/H_sapiens/annotation/GRCh37_latest/refseq_identifiers/GRCh37_latest_genomic.gff.gz |
| Accessibility masks | http://ftp.1000genomes.ebi.ac.uk/vol1/ftp/release/20130502/supporting/accessible_genome_masks/20140520.strict_mask.autosomes.bed |
| Phastcons masks | http://hgdownload.soe.ucsc.edu/goldenPath/hg19/phastCons100way/hg19.100way.phastCons.bw |
| Recombination rate data | https://hgdownload.soe.ucsc.edu/gbdb/hg19/decode/SexAveraged.bw |
| Mutation rate data | https://download.molgeniscloud.org/downloads/gonl_public/mutation_rate_map/release2/local_mutation_rate.bias_corrected.SEXAVG.bed |
| GRCh37 variation data | http://ftp.1000genomes.ebi.ac.uk/vol1/ftp/release/20130502/ |
| GRCh37 reference | Obtained from Enseml rest API |
| Ancestral sequences | https://ftp.ensembl.org/pub/release-74/fasta/ancestral_alleles/homo_sapiens_ancestor_GRCh37_e71.tar.bz2 |

| Parameter | Parameter range | | Units | Notes | Citation |
|---|---|---|---|---|---|
| | Lower | Upper | | | |
| $\tau_{AFR\text{-}EURASI}$ | 45 | 140 | kya | $\tau_{X\text{-}Y}$ is split time of population Y from population X | Henn et al. 2012 |
| $\tau_{EURASI\text{-}EUR}$ | 17.2 | 40 | | | Gutenkunst et al. 2009; Mellars 2006 |
| $\tau_{EURASI\text{-}EAS}$ | 17.2 | 40 | | | Gutenkunst et al. 2009; Mellars 2006 |
| $\tau_{EURASI\text{-}SAS}$ | 17.2 | 40 | | | Gutenkunst et al. 2009; Mellars 2006 |
| $\tau_{AFR}$ | 5 | 15 | | | Tennessen et al. 2012; Terhorst et al. 2017 |
| $N_{AFRancestral}$ | 4,400 | 14,400 | $N$ | Ancestral African population size | Gutenkunst et al. 2009; Henn et al. 2012 |
| $B_{EURASI}$<br>$B_{EUR}$<br>$B_{EAS}$<br>$B_{SAS}$ | 0.01 | 0.2 | $N$ | Population size at dispersal from ancestral population via bottleneck ranging between 99% and 80% severity. | n/a |

| | | | | | |
|---|---|---|---|---|---|
| $r_{AFR}$<br>$r_{EURASI}$<br>$r_{EUR}$<br>$r_{EAS}$<br>$r_{SAS}$ | 0.1 | 1 | % | Growth rate per generation | Gutenkunst et al. 2009; Gravel et al. 2006 |
| $m_{AFR-EURASI}$<br>$m_{AFR-EUR}$<br>$m_{AFR-EAS}$<br>$m_{AFR-SAS}$<br>$m_{EURASI-EUR}$<br>$m_{EURASI-EAS}$<br>$m_{EURASI-SAS}$<br>$m_{EUR-EAS}$<br>$m_{EUR-SAS}$<br>$m_{EAS-SAS}$ | 0 | 10 | $4Nm$ | $m_{X-Y}$ is the symmetrical migration rate between population X and population Y. Range between $4Nm$=0 (isolation) to $4Nm$=10 (panmixia). | n/a |

**Table S2:** Tested parameter ranges for human Out-Of-Africa demographic model

**S1**

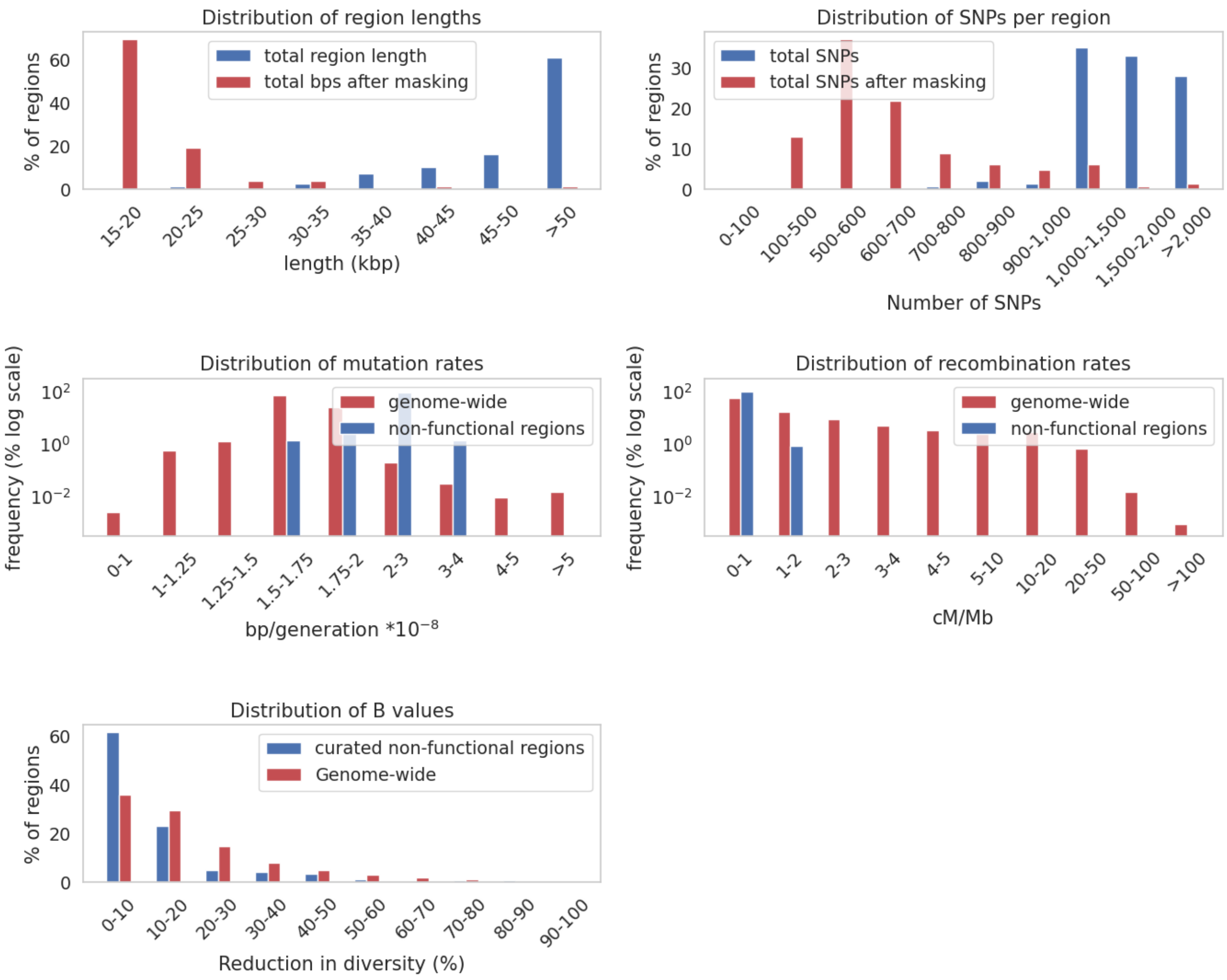


**Supplementary Figure S1:** Upper Left: Distribution of lengths of curated non-functional regions after accessibility and phastCons masking. Upper right: Distribution of SNPs per region. Middle left: Comparison of distributions of mutation rates genome-wide and for curated non-functional regions. Genome-wide rates calculated over 20kb windows. Middle right: Comparison of distributions of recombination rates genome-wide and for curated non-functional regions. Bottom left: Comparison of distribution of genome-wide B values with B values across our 146 curated non-functional regions.

**S2**

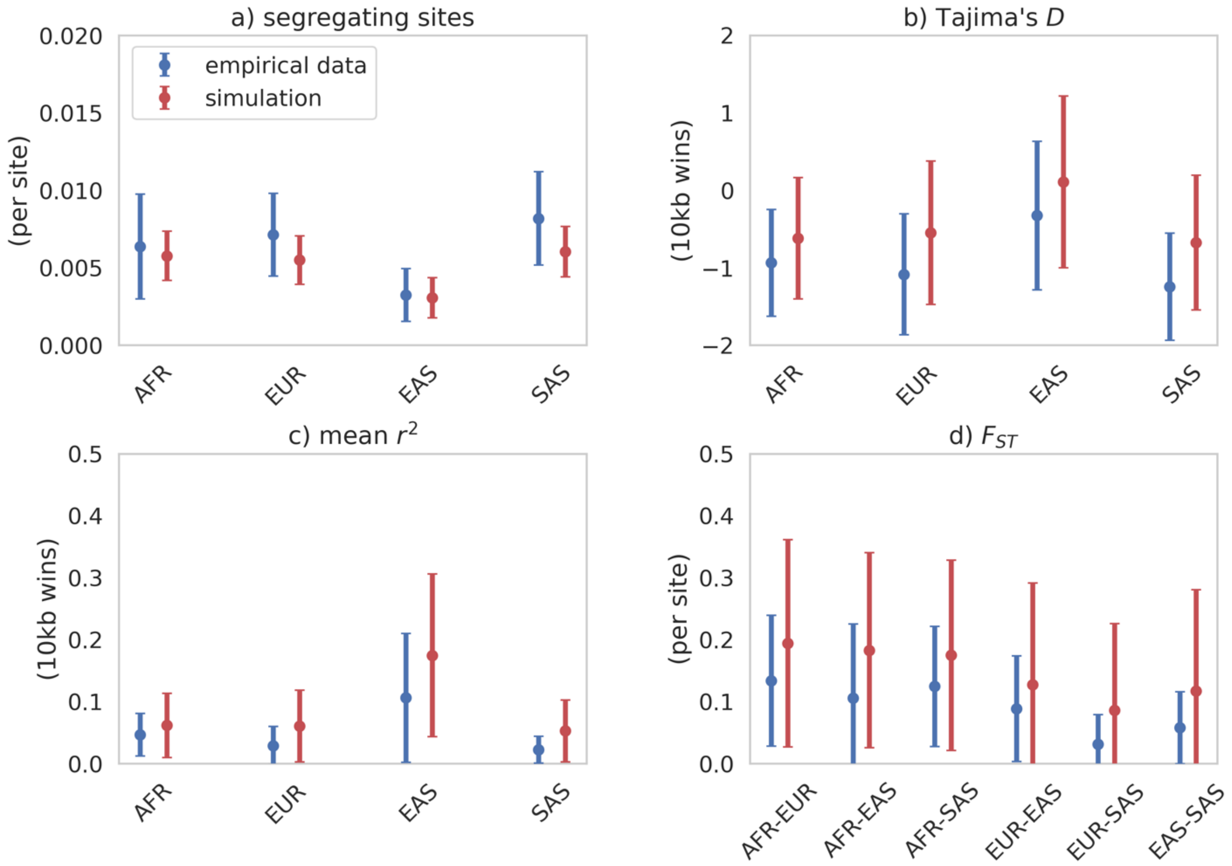


**Supplementary Figure S2:** Summary statistics calculated from functional regions from each sampled population for empirical and simulated data. Simulated data comes from our best-fitting neutral demographic model parametrizations (see Figure 1). Means and standard deviations were calculated across 100 replicates. Data points represent the means across regions, while bars represent the mean of the standard deviations across regions.

**S3**

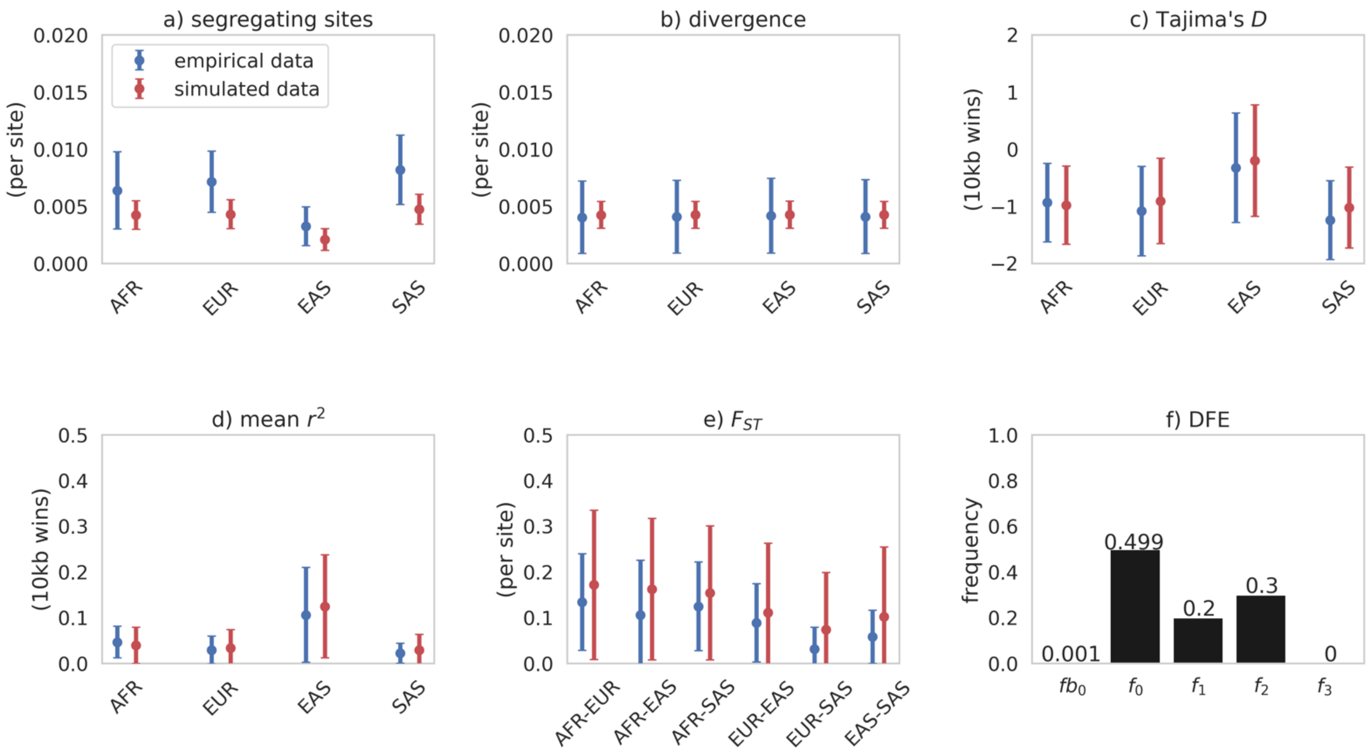


**Supplementary Figure S3:** a) to e) Summary statistics calculated from functional regions for each sampled population for empirical and simulated data. Simulated data comes from our best-fitting demographic model parametrizations (see Figure 1), with purifying selection and BGS modelled using the Johri et al. (2023) DFE, except with the addition of a beneficial mutational category (panel f). Means and standard deviations were calculated across 100 replicates. Data points represent the means across regions, while bars represent the mean of the standard deviations across regions. Exonic mutations were drawn from a DFE comprised of 5 fixed classes with frequencies denoted by $f_i$ and $f_{b0}$: $f_{b0}$ with $1 \leq 2N_{AFRancestral}\, s_b < 10$ (i.e., weakly beneficial mutations), $f_0$ with $0 \leq 2N_{AFRancestral}\, s < 1$ (i.e., effectively neutral mutations), $f_1$ with $1 \leq 2N_{AFRancestral}\, s < 10$ (i.e., weakly deleterious mutations), $f_2$ with $10 \leq 2N_{AFRancestral}\, s < 100$ (i.e., moderately deleterious mutations), and $f_3$ with $100 \leq 2N_{AFRancestral}\, s$ (i.e., strongly deleterious mutations), where $N_{AFRancestral}$ was the initial population size, $s$ the reduction in fitness of the mutant homozygote relative to wild-type, and $s_b$ the increase in fitness of the beneficial mutation. **Here, weakly beneficial mutations comprise 0.1% of new mutations.**

**S4**

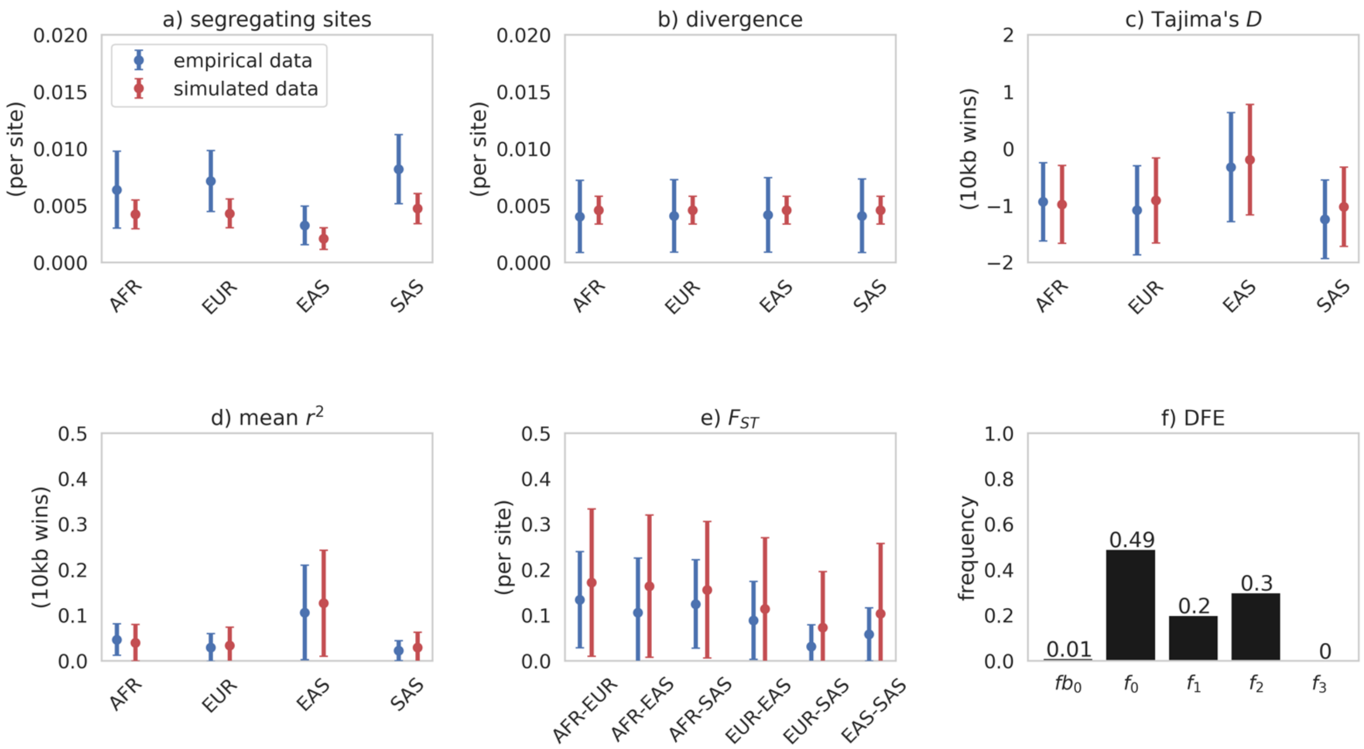


**Supplementary Figure S4:** a) to e) Summary statistics calculated from functional regions for each sampled population for empirical and simulated data. Simulated data comes from our best-fitting demographic model parametrizations (see Figure 1), with purifying selection and BGS modelled using the Johri et al. (2023) DFE, except with the addition of a beneficial mutational category (panel f). Means and standard deviations were calculated across 100 replicates. Data points represent the means across regions, while bars represent the mean of the standard deviations across regions. Exonic mutations were drawn from a DFE comprised of 5 fixed classes with frequencies denoted by $f_i$ and $f_{b0}$: $f_{b0}$ with $1 \leq 2N_{AFRancestral}\, s_b < 10$ (i.e., weakly beneficial mutations), $f_0$ with $0 \leq 2N_{AFRancestral}\, s < 1$ (i.e., effectively neutral mutations), $f_1$ with $1 \leq 2N_{AFRancestral}\, s < 10$ (i.e., weakly deleterious mutations), $f_2$ with $10 \leq 2N_{AFRancestral}\, s < 100$ (i.e., moderately deleterious mutations), and $f_3$ with $100 \leq 2N_{AFRancestral}\, s$ (i.e., strongly deleterious mutations), where $N_{AFRancestral}$ was the initial population size, $s$ the reduction in fitness of the mutant homozygote relative to wild-type, and $s_b$ the increase in fitness of the beneficial mutation. **Here, weakly beneficial mutations comprise 1% of new mutations.**

**S5**

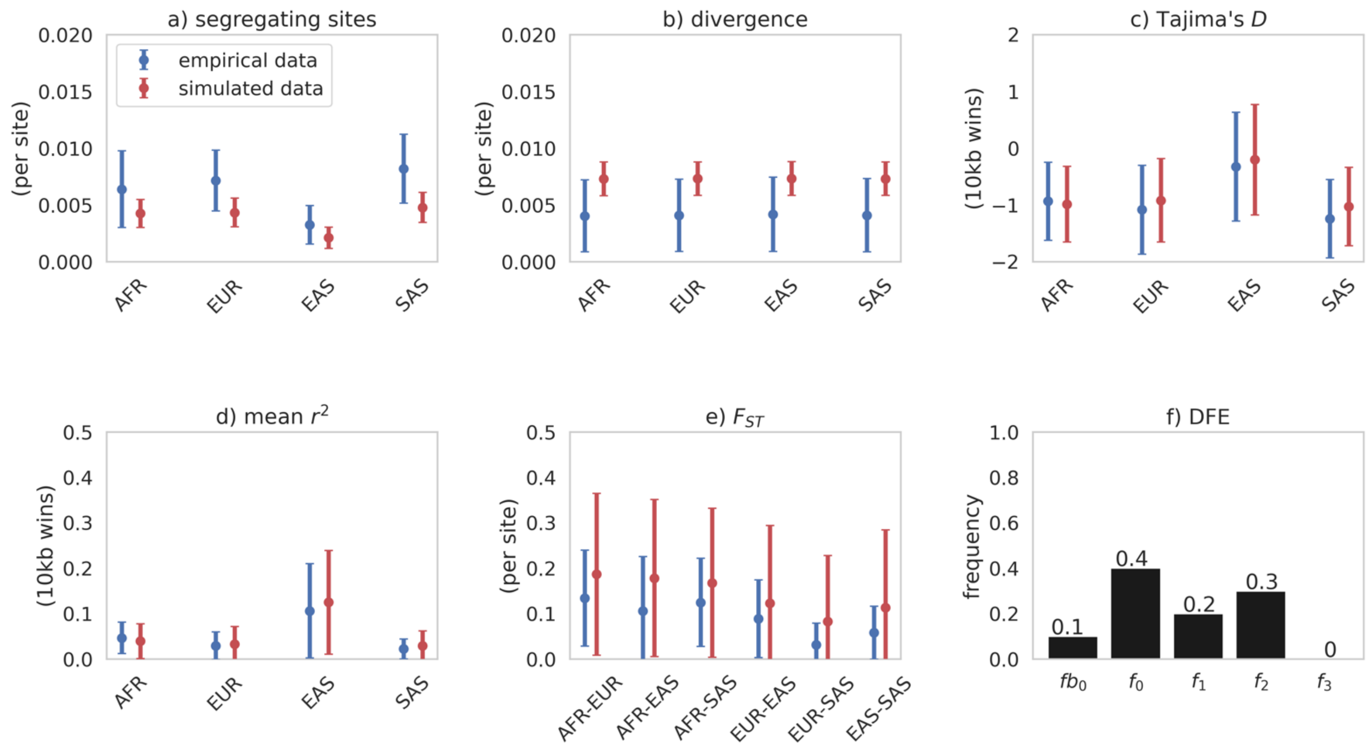


**Supplementary Figure S5:** a) to e) Summary statistics calculated from functional regions for each sampled population for empirical and simulated data. Simulated data comes from our best-fitting demographic model parametrizations (see Figure 1), with purifying selection and BGS modelled using the Johri et al. (2023) DFE, except with the addition of a beneficial mutational category (panel f). Means and standard deviations were calculated across 100 replicates. Data points represent the means across regions, while bars represent the mean of the standard deviations across regions. Exonic mutations were drawn from a DFE comprised of 5 fixed classes with frequencies denoted by $f_i$ and $f_{b0}$: $f_{b0}$ with $1 \leq 2N_{AFRancestral}\, s_b < 10$ (i.e., weakly beneficial mutations), $f_0$ with $0 \leq 2N_{AFRancestral}\, s < 1$ (i.e., effectively neutral mutations), $f_1$ with $1 \leq 2N_{AFRancestral}\, s < 10$ (i.e., weakly deleterious mutations), $f_2$ with $10 \leq 2N_{AFRancestral}\, s < 100$ (i.e., moderately deleterious mutations), and $f_3$ with $100 \leq 2N_{AFRancestral}\, s$ (i.e., strongly deleterious mutations), where $N_{AFRancestral}$ was the initial population size, $s$ the reduction in fitness of the mutant homozygote relative to wild-type, and $s_b$ the increase in fitness of the beneficial mutation. **Here, weakly beneficial mutations comprise 10% of new mutations.**

**S6**

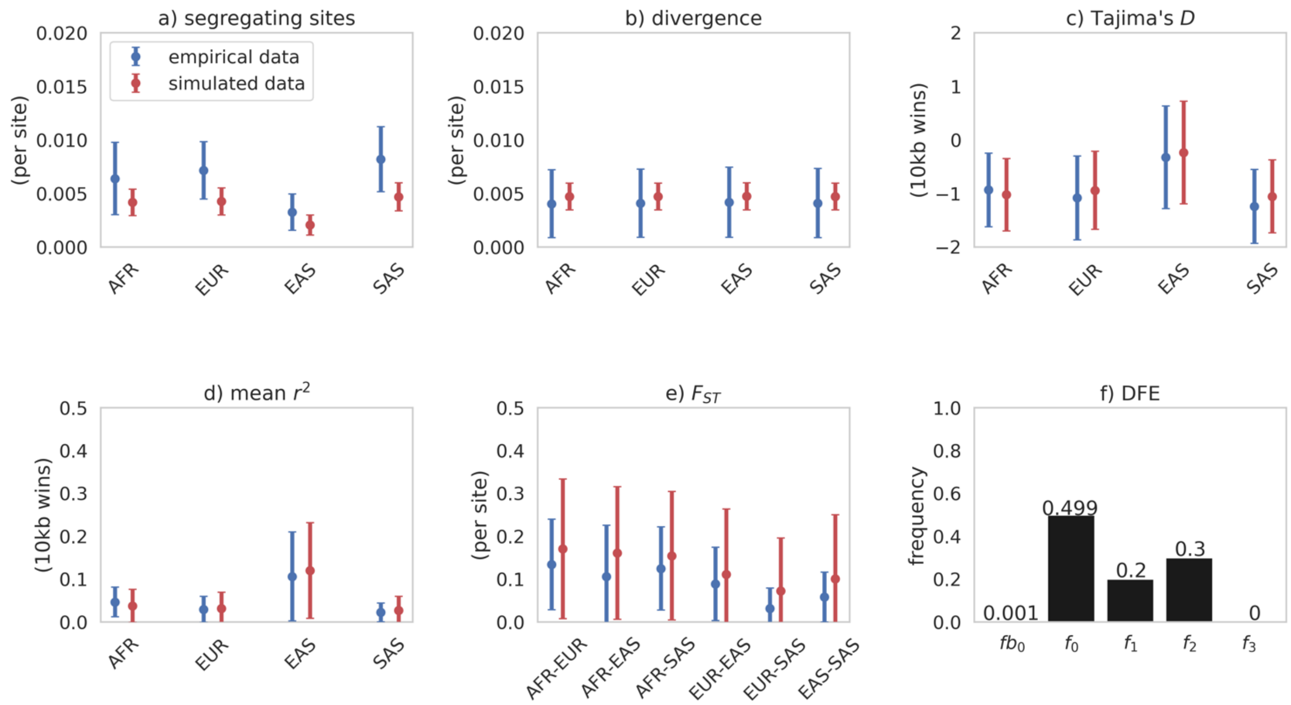


**Supplementary Figure S6:** a) to e) Summary statistics calculated from functional regions for each sampled population for empirical and simulated data. Simulated data comes from our best-fitting demographic model parametrizations (see Figure 1), with purifying selection and BGS modelled using the Johri et al. (2023) DFE, except with the addition of a beneficial mutational category (panel f). Means and standard deviations were calculated across 100 replicates. Data points represent the means across regions, while bars represent the mean of the standard deviations across regions. Exonic mutations were drawn from a DFE comprised of 5 fixed classes with frequencies denoted by $f_i$ and $f_{b0}$: $f_{b0}$ with $10 \leq 2N_{AFRancestral}$ $s_b < 100$ (i.e., moderately beneficial mutations), $f_0$ with $0 \leq 2N_{AFRancestral}\ s < 1$ (i.e., effectively neutral mutations), $f_1$ with $1 \leq 2N_{AFRancestral}\ s < 10$ (i.e., weakly deleterious mutations), $f_2$ with $10 \leq 2N_{AFRancestral}l\ s < 100$ (i.e., moderately deleterious mutations), and $f_3$ with $100 \leq$ $2N_{AFRancestral}\ s$ (i.e., strongly deleterious mutations), where $N_{AFRancestral}$ was the initial population size, $s$ the reduction in fitness of the mutant homozygote relative to wild-type, and $s_b$ the increase in fitness of the beneficial mutation. **Here, moderately beneficial mutations comprise 0.1% of new mutations.**

**S7**

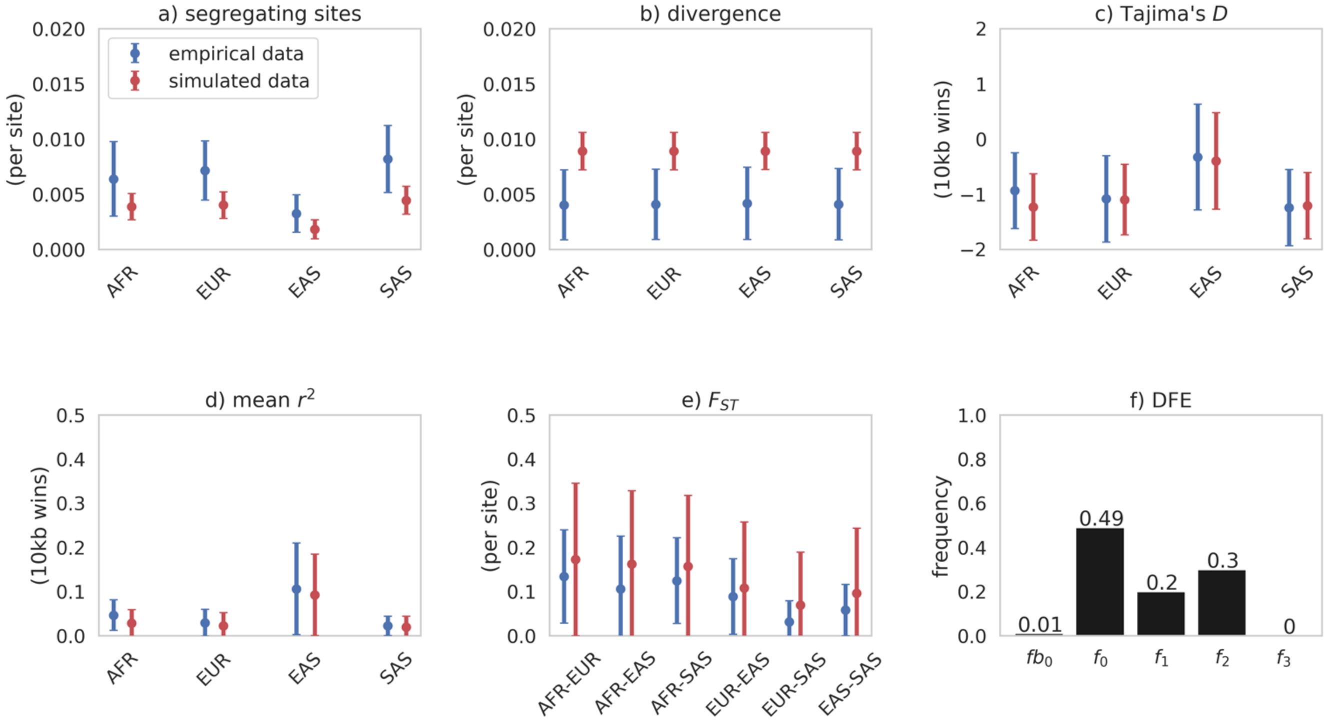


**Supplementary Figure S7:** a) to e) Summary statistics calculated from functional regions for each sampled population for empirical and simulated data. Simulated data comes from our best-fitting demographic model parametrizations (see Figure 1), with purifying selection and BGS modelled using the Johri et al. (2023) DFE, except with the addition of a beneficial mutational category (panel f). Means and standard deviations were calculated across 100 replicates. Data points represent the means across regions, while bars represent the mean of the standard deviations across regions. Exonic mutations were drawn from a DFE comprised of 5 fixed classes with frequencies denoted by $f_i$ and $f_{b0}$: $f_{b0}$ with $10 \le 2N_{AFRancestral}\, s_b < 100$ (i.e., moderately beneficial mutations), $f_0$ with $0 \le 2N_{AFRancestral}\, s < 1$ (i.e., effectively neutral mutations), $f_1$ with $1 \le 2N_{AFRancestral}\, s < 10$ (i.e., weakly deleterious mutations), $f_2$ with $10 \le 2N_{AFRancestral}\, s < 100$ (i.e., moderately deleterious mutations), and $f_3$ with $100 \le 2N_{AFRancestral}\, s$ (i.e., strongly deleterious mutations), where $N_{AFRancestral}$ was the initial population size, $s$ the reduction in fitness of the mutant homozygote relative to wild-type, and $s_b$ the increase in fitness of the beneficial mutation. **Here, moderately beneficial mutations comprise 1% of new mutations.**

**S8**

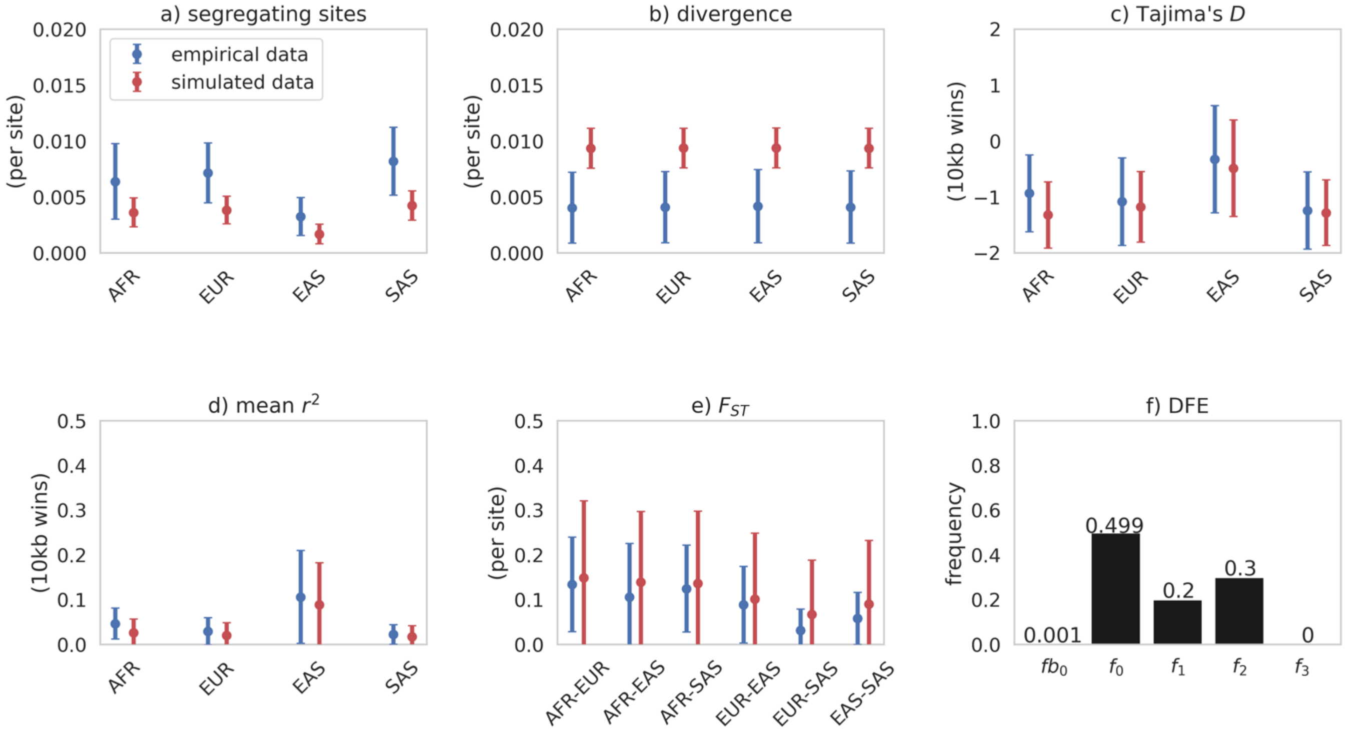


**Supplementary Figure S8:** a) to e) Summary statistics calculated from functional regions for each sampled population for empirical and simulated data. Simulated data comes from our best-fitting demographic model parametrizations (see Figure 1), with purifying selection and BGS modelled using the Johri et al. (2023) DFE, except with the addition of a beneficial mutational category (panel f). Means and standard deviations were calculated across 100 replicates. Data points represent the means across regions, while bars represent the mean of the standard deviations across regions. Exonic mutations were drawn from a DFE comprised of 5 fixed classes with frequencies denoted by $f_i$ and $f_{b0}$: $f_{b0}$ with $100 \leq 2N_{AFRancestral}\ s_b < 1000$ (i.e., strongly beneficial mutations), $f_0$ with $0 \leq 2N_{AFRancestral}\ s < 1$ (i.e., effectively neutral mutations), $f_1$ with $1 \leq 2N_{AFRancestral}\ s < 10$ (i.e., weakly deleterious mutations), $f_2$ with $10 \leq 2N_{AFRancestral}\ s < 100$ (i.e., moderately deleterious mutations), and $f_3$ with $100 \leq 2N_{AFRancestral}\ s$ (i.e., strongly deleterious mutations), where $N_{AFRancestral}$ was the initial population size, $s$ the reduction in fitness of the mutant homozygote relative to wild-type, and $s_b$ the increase in fitness of the beneficial mutation. **Here, strongly beneficial mutations comprise 0.1% of new mutations.**

**S9**

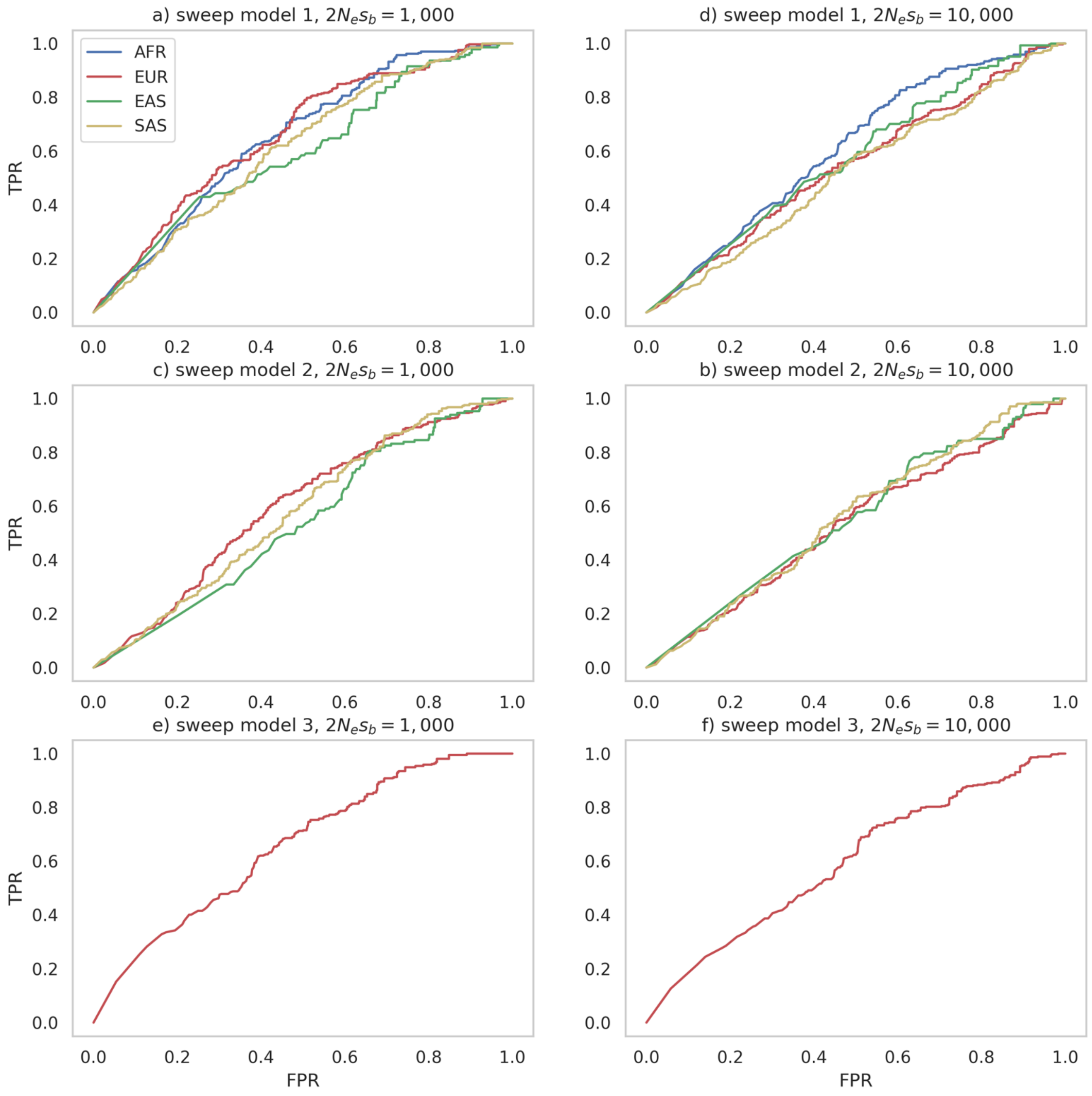


**Supplementary Figure S9:** ROC curves, providing the change in true-positive rate (TPR) with false-positive rate (FPR), for sweep inference with the H12 statistic, under the OOA demographic model inferred in this study (see Figure 1), together with the Johri et al. (2023) DFE for functional regions with the addition of single beneficial mutation, as well as variable mutation and recombination rates (see Methods). The beneficial mutation was introduced into the simulated population at three different times and in three different populations: Model 1: the beneficial mutation was introduced into the ancestral African population immediately after the burn-in period, the fixed beneficial mutation is present in all populations, and sweep inference was conducted on all populations; Model 2: the beneficial mutation was introduced into the ancestral Eurasian population immediately upon splitting

from the African population, the fixed beneficial mutation is present in all non-African populations, and sweep inference was conducted on the European, East Asian and South Asian populations; Model 3: the beneficial mutation was introduced into the European population immediately upon splitting from the Eurasian population, the beneficial mutation is fixed in the European population, and sweep inference was conducted on this population only. For each model, two different strengths of selection were considered: $2N_es_b$=1,000 and $2N_es_b$=10,000, where $N_e$ is the size of the ancestral African population and $s_b$ is the beneficial selection coefficient. Inference was performed on each SNP over a **2kb window**, with the SNP at the center of the window.

**S10**

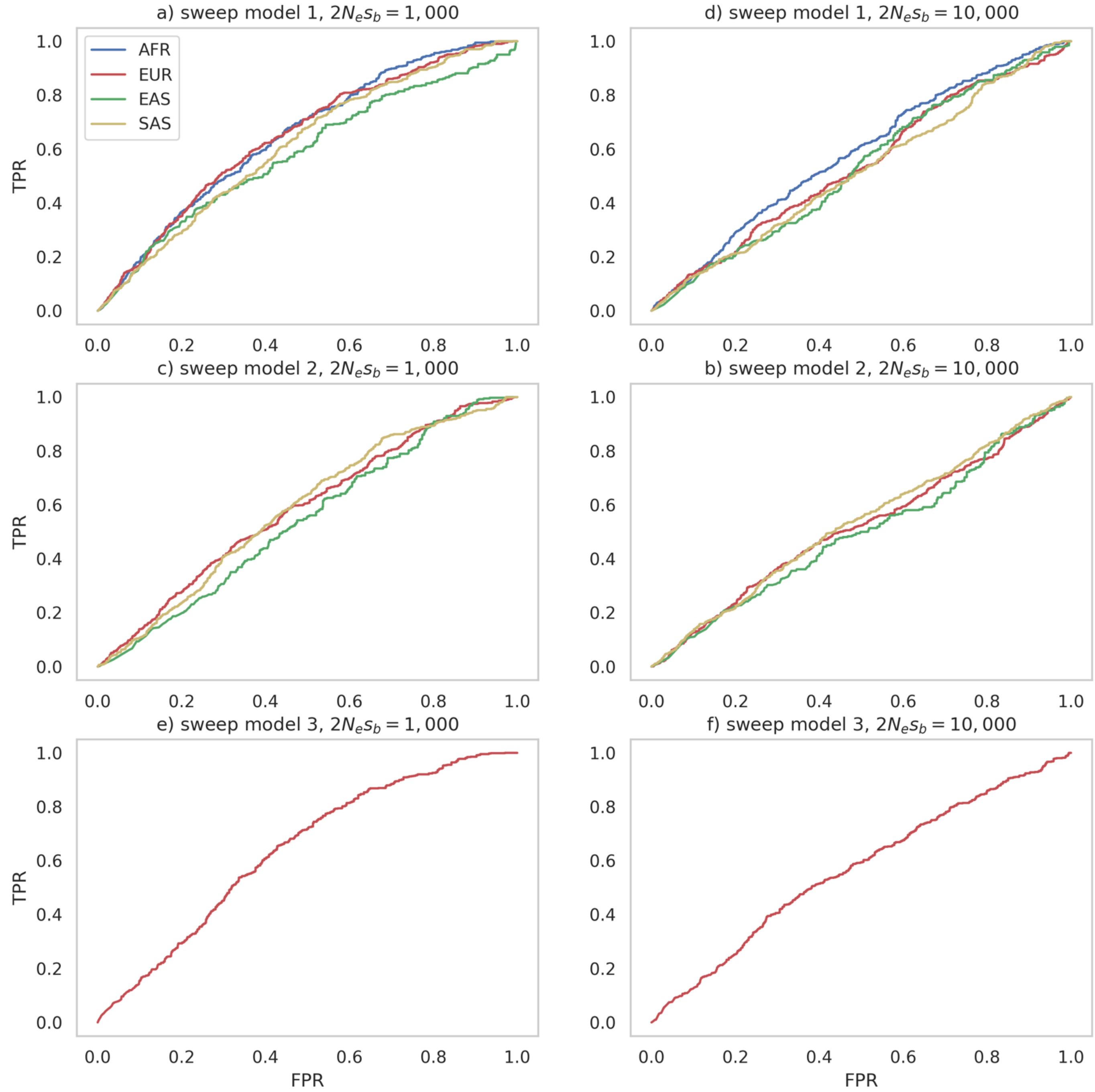


**Supplementary Figure S10:** ROC curves, providing the change in true-positive rate (TPR) with false-positive rate (FPR), for sweep inference with the H12 statistic, under the OOA demographic model inferred in this study (see Figure 1), together with the Johri et al. (2023) DFE for functional regions with the addition of single beneficial mutation, as well as variable mutation and recombination rates (see Methods). The beneficial mutation was introduced into the simulated population at three different times and in three different populations: Model 1: the beneficial mutation was introduced into the ancestral African population immediately after the burn-in period, the fixed beneficial mutation is present in all populations, and sweep inference was conducted on all populations; Model 2: the beneficial mutation was introduced into the ancestral Eurasian population immediately upon splitting

from the African population, the fixed beneficial mutation is present in all non-African populations, and sweep inference was conducted on the European, East Asian and South Asian populations; Model 3: the beneficial mutation was introduced into the European population immediately upon splitting from the Eurasian population, the beneficial mutation is fixed in the European population, and sweep inference was conducted on this population only.  For each model, two different strengths of selection were considered: $2N_es_b$=1,000 and $2N_es_b$=10,000, where $N_e$ is the size of the ancestral African population and $s_b$ is the beneficial selection coefficient. Inference was performed on each SNP over a **5kb window**, with the SNP at the center of the window.

**S11**

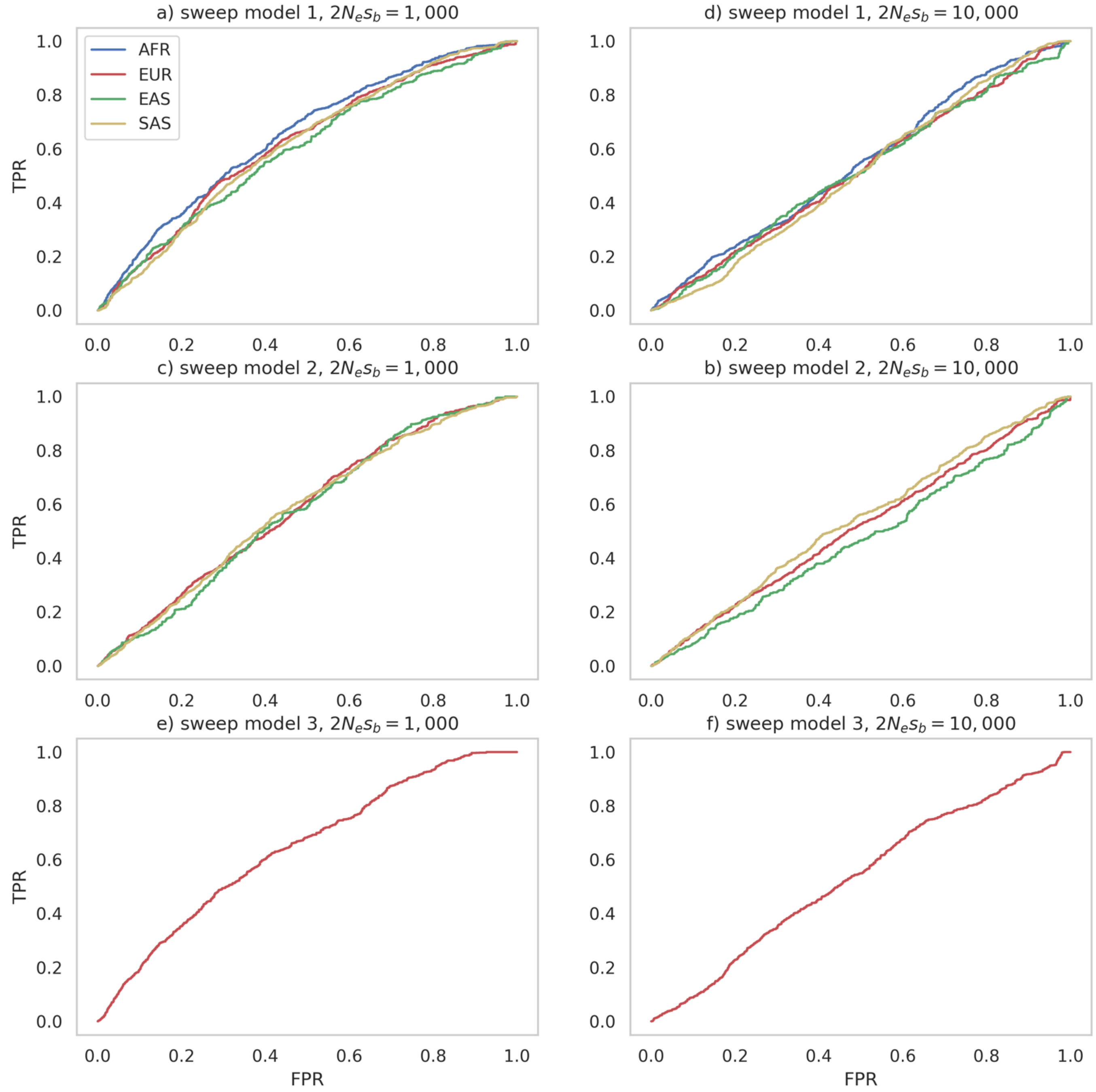


**Supplementary Figure S11:** ROC curves, providing the change in true-positive rate (TPR) with false-positive rate (FPR), for sweep inference with the H12 statistic, under the OOA demographic model inferred in this study (see Figure 1), together with the Johri et al. (2023) DFE for functional regions with the addition of single beneficial mutation, as well as variable mutation and recombination rates (see Methods). The beneficial mutation was introduced into the simulated population at three different times and in three different populations: Model 1: the beneficial mutation was introduced into the ancestral African population immediately after the burn-in period, the fixed beneficial mutation is present in all populations, and sweep inference was conducted on all populations; Model 2: the beneficial mutation was introduced into the ancestral Eurasian population immediately upon splitting

from the African population, the fixed beneficial mutation is present in all non-African populations, and sweep inference was conducted on the European, East Asian and South Asian populations; Model 3: the beneficial mutation was introduced into the European population immediately upon splitting from the Eurasian population, the beneficial mutation is fixed in the European population, and sweep inference was conducted on this population only.  For each model, two different strengths of selection were considered: $2N_es_b$=1,000 and $2N_es_b$=10,000, where $N_e$ is the size of the ancestral African population and $s_b$ is the beneficial selection coefficient. Inference was performed on each SNP over a **10kb window**, with the SNP at the center of the window.

**S12**

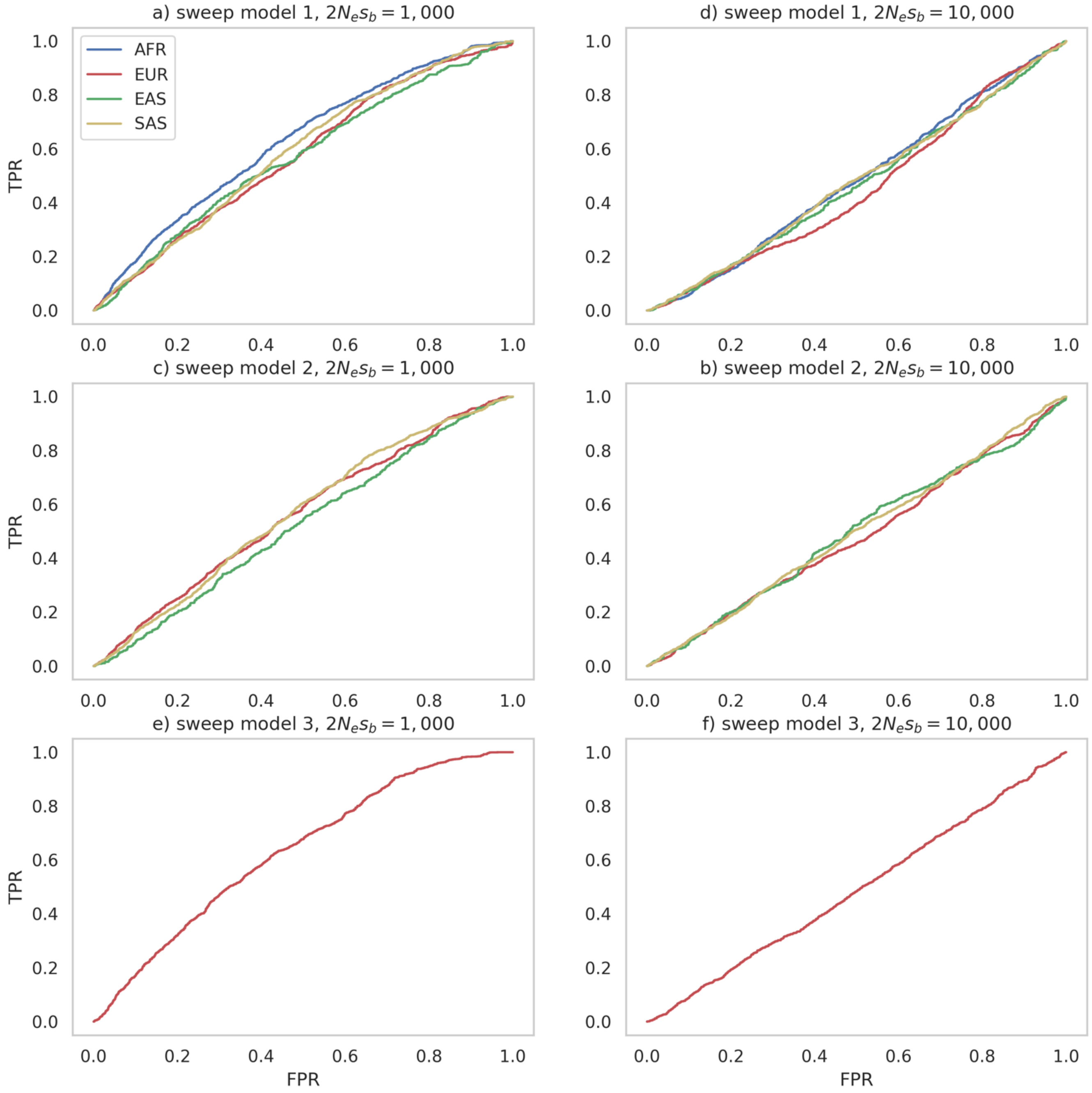


**Supplementary Figure S12:** ROC curves, providing the change in true-positive rate (TPR) with false-positive rate (FPR), for sweep inference with the H12 statistic, under the OOA demographic model inferred in this study (see Figure 1), together with the Johri et al. (2023) DFE for functional regions with the addition of single beneficial mutation, as well as variable mutation and recombination rates (see Methods). The beneficial mutation was introduced into the simulated population at three different times and in three different populations: Model 1: the beneficial mutation was introduced into the ancestral African population immediately after the burn-in period, the fixed beneficial mutation is present in all populations, and sweep inference was conducted on all populations; Model 2: the beneficial mutation was introduced into the ancestral Eurasian population immediately upon splitting

from the African population, the fixed beneficial mutation is present in all non-African populations, and sweep inference was conducted on the European, East Asian and South Asian populations; Model 3: the beneficial mutation was introduced into the European population immediately upon splitting from the Eurasian population, the beneficial mutation is fixed in the European population, and sweep inference was conducted on this population only. For each model, two different strengths of selection were considered: $2N_es_b$=1,000 and $2N_es_b$=10,000, where $N_e$ is the size of the ancestral African population and $s_b$ is the beneficial selection coefficient. Inference was performed on each SNP over a **20kb window**, with the SNP at the center of the window.

**S13**

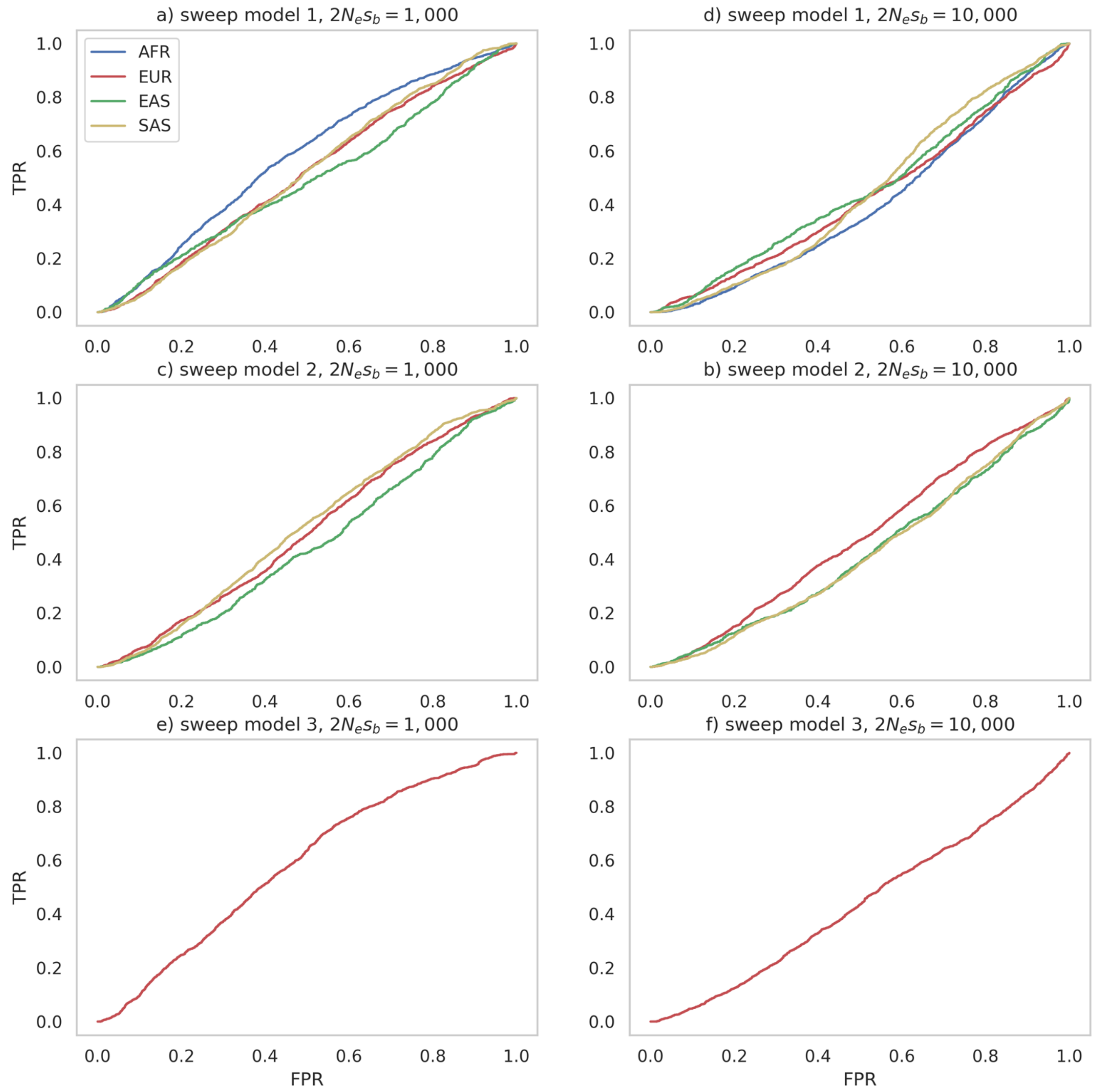


**Supplementary Figure S13:** ROC curves, providing the change in true-positive rate (TPR) with false-positive rate (FPR), for sweep inference with the H12 statistic, under the OOA demographic model inferred in this study (see Figure 1), together with the Johri et al. (2023) DFE for functional regions with the addition of single beneficial mutation, as well as variable mutation and recombination rates (see Methods). The beneficial mutation was introduced into the simulated population at three different times and in three different populations: Model 1: the beneficial mutation was introduced into the ancestral African population immediately after the burn-in period, the fixed beneficial mutation is present in all populations, and sweep inference was conducted on all populations; Model 2: the beneficial mutation was introduced into the ancestral Eurasian population immediately upon splitting from the African population, the fixed beneficial mutation is present in all non-African populations, and sweep inference was conducted on the European, East Asian and South Asian populations; Model 3: the beneficial mutation was introduced into the European population immediately upon splitting from the Eurasian population, the beneficial mutation is fixed in the European population, and sweep inference was conducted on this population only. For each model, two different strengths of selection were considered: $2N_es_b$=1,000 and $2N_es_b$=10,000, where $N_e$ is the size of the ancestral African population and $s_b$ is the beneficial selection coefficient. Inference was performed on each SNP over a **40kb window**, with the SNP at the center of the window.
